# Nonstoichiometric balanced complexes: Implications on the effective deficiency of the underlying metabolic network

**DOI:** 10.1101/2021.07.07.451418

**Authors:** Damoun Langary, Anika Küken, Zoran Nikoloski

**Affiliations:** Bioinformatics, Institute of Biochemistry and Biology, University of Potsdam, Potsdam, Germany; Systems Biology and Mathematical Modeling, Max Planck Institute of Molecular Plant Physiology, Potsdam, Germany

**Keywords:** balanced complexes, constraint-based modeling, chemical reaction network theory, deficiency, metabolic networks

## Abstract

The deficiency of a (bio)chemical reaction network can be conceptually interpreted as a measure of its ability to support exotic dynamical behavior and/or multistationarity. The classical definition of deficiency relates to the capacity of a network to permit variations of the complex formation rate vector at steady state, irrespective of the network kinetics. However, the deficiency is by definition completely insensitive to the fine details of the directionality of reactions as well as bounds on reaction fluxes. While the classical definition of deficiency can be readily applied in the analysis of unconstrained, weakly reversible networks, it only provides an upper bound in the cases where relevant constraints on reaction fluxes are imposed. Here we propose the concept of effective deficiency, which provides a more accurate assessment of the network’s capacity to permit steady state variations at the complex level for constrained networks of any reversibility patterns. The effective deficiency relies on the concept of nonstoichiometric balanced complexes, which we have already shown to be present in real-world biochemical networks operating under flux constraints. Our results demonstrate that the effective deficiency of real-world biochemical networks is smaller than the classical deficiency, indicating the effects of reaction directionality and flux bounds on the variation of the complex formation rate vector at steady state.

## 1 Introduction

Chemical reaction network theory (CRNT) has made seminal contributions to establishing necessary and/or sufficient conditions that a (bio)chemical network exhibits particular properties, such as: presence / absence of multiple steady state, stability of steady states, and robustness of steady-state concentrations (Feinberg, 2019). The deficiency of a network is of central importance in this theory, and many steady-state concentration properties are guaranteed or precluded for networks of particular deficiency (Feinberg, 1987; 1988; Shinar & Feinberg, 2011). These results often hold for (bio)chemical networks whose reactions are endowed with (generalized) mass action kinetics. However, a notable feature of this theory is that it does not impose any physico-chemical constraints that real-world networks obey. As a result, the notion of deficiency is not concerned with the (ir)reversibility of the considered biochemical reactions that may arise in practice, and the extent to which this may affect the properties of the corresponding steady states the network supports.

In contrast to CRNT, which almost exclusively deals with concentration-based properties, the constraint-based modeling framework (Bordbar, et al., 2014; Orth, et al., 2010), imposes physico-chemical constraints as lower and/or upper bounds on reaction rates (i.e. fluxes) in making predictions about macro-level phenotypes, such as growth or yield of a chemical product of interest. The constraint-based modeling framework has found numerous applications, largely due to its capacity to model large-scale networks by building on approaches for convex optimization.

A natural question then arises of how to integrate any imposed constraints on reaction fluxes into the notion of deficiency. In addition one can ask whether or not a given (bio)chemical network is of lower deficiency compared to that of the classical definition, solely based on the network structure and imposed bounds. In such a way, one may expand the usability of the classical results from CRNT in networks of seemingly different class, based on the classical notion of deficiency (Feinberg, 1987; 1988). Work in this direction has led to the concept of network translation that allows to cast a (bio)chemical network endowed with generalized mass action kinetics by a network of lower deficiency and other structural properties (Johnston, 2014).

Here, we rely on the recently introduced classification of balanced complexes (Langary, et al., 2021), in particular the class of nonstoichiometric balanced complexes, to define the notion of effective deficiency of a network. Nonstoichiometric balanced complexes have been shown to arise as a combined result of several factors, including the algebraic and graphical structure of the network as well as operational bounds on reaction kinetics (Langary, et al., 2021). A remarkable feature of the effective deficiency is that, like that of the classical, so-called structural, it is defined and can be calculated for networks of arbitrary kinetics.

Our theoretical results show that decrease in the structural deficiency of a network is due to the presence of reactions whose fluxes are fixated at their lower bounds. In other words, given the irreversibility and flux bound constraints, if the network contains nonstoichiometric balanced complexes, some reactions are guaranteed to exhibit absolute flux robustness. It is then not surprising that the effective deficiency captures the reduced capacity for variation of the complex formation rate at steady state.

Finally, application of our results on twelve real-world metabolic networks of species from all three kingdoms of life show that their effective deficiency is decreased by 8% on average in comparison to the structural deficiency, with the highest decrease in deficiency of 35% observed for the network of *M. barkeri.* These results along with those from selected illustrative toy networks demonstrate the implications of our results to precluding exotic dynamic behavior asserted by well-established results based on the classical notion of structural deficiency.

## 2 Background on CRNT

### 2.1 Problem Setup

A chemical reaction network (CRN) is defined by a set of *m species/metabolites*, 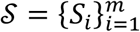, a set of *n complexes*, 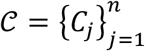 each of which is a multiset of species 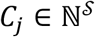, and a set of *r reactions*, 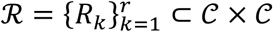 which symbolize the potential conversion of complexes into each other in the network (Horn & Jackson, 1972; Feinberg, 1980; 1987; 1988).

We denote the standard basis in 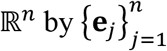 where

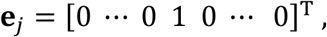

is an *n*-vector with a unit value at the *j*^th^ entry and zero entries elsewhere. Let us next assume some arbitrary ordering on the sets of species (*S*_1_, …, *S_m_*), complexes (*C*_1_, …, *C_n_*), and reactions (*R*_1_, …, *R_r_*). Given this ordering, any complex 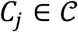 can be associated with a vector 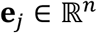 indexing its position in the ordered set, and at the same time with a unique vector 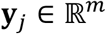, which represents its species content. This defines the following mapping

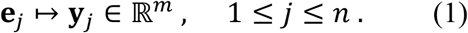

Consequently, any given CRN is associated with a *stoichiometric map*, defined by the matrix 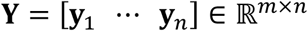. Similarly, the ordering associates each reaction 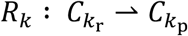 with a vector 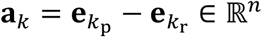 representing the conversion in the complex space, and a vector 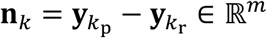, representing the conversion in the species space. Thus, the CRN is at the same time associated with the *complex - reaction incidence matrix* 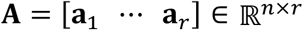 and a species conversion matrix 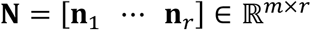, known as *stoichiometry matrix.* It follows immediately from the definition of the stoichiometric map that **N** = **Y A**. The column span of **N** is called the *stoichiometric subspace*, the dimension of which, *s*, is termed *rank* of the CRN, that is, *s* = rank(**N**).

Assuming the reaction network is endowed with some kinetics, at any state of the system, denoted by the vector of concentrations 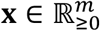, the dynamics is governed by the following ODE

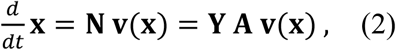

where 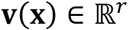 returns the reaction rates determined by system kinetics as a generally nonlinear function of the state vector *x*.

### 2.2 Blocked reactions and the steady state flux space

The vector 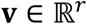, referred to as a *flux distribution* for the network 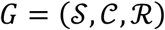, is said to be at steady state, if

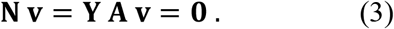

In addition, the flux vector **v** is often assumed to be bounded by box constraints as follows

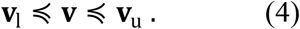

where 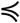 operates element-wise, and 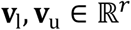 specify the individual lower- and upper bounds on flux through reactions, respectively. By convention, **v**_u_ is strictly positive for all reactions, 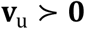. Going forward, we assume a network denoted *G* also encodes information about upper and lower bounds on reaction fluxes.

When no additional restriction is imposed on the network, a flux distribution **v** is called *feasible*, if it satisfies both constraints (3) and (4). The set of all feasible flux distributions is an intercept of some hyperplanes and halfspaces, which defines a convex polyhedron, referred to as the *steady state flux space*, denoted 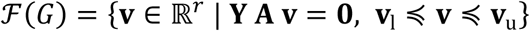.

The set of reversible reactions 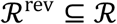 is defined by negative entries of **V**_l_, that is

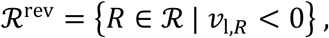

where *υ*_l,*R*_ denotes the entry of **v**_l_ corresponding to reaction *R*. The set of irreversible reactions 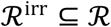 is then defined as the complement of 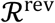, that is, 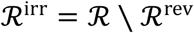. Hence, irreversible reactions are associated with *υ*_l,*R*_ ≥ 0. By contrast to irreversible reactions, a reversible reaction 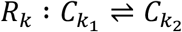 portrays the mutual conversion of complexes *C*_*k*_1__ and *C*_*k*_2__.

Similarly, let 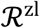 and 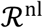 denote the sets of reactions with zero and nonzero lower bounds, respectively. It follows from the definition that, 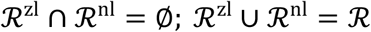; and 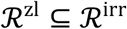. Given the arbitrariness of indexing, one can always order the reactions in 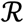 such that elements of 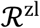 and 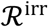 precede those of 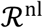 and 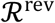, respectively. As a result, the incidence matrix **A** can be block-partitioned as either **A** = [**A**^irr^ **A**^rev^] or **A** = [**A**^zl^ **A**^nl^].

The network is said to be operating under a *canonical flux regime*, if 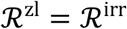. It is said to be operating under an *unbounded flux regime*, if **v**_u_ = ∞ and *υ*_l,*R*_ = –∞, 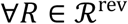. Under a canonical and unbounded flux regime, the set of feasible flux distributions forms a polyhedral convex cone, referred to as the *steady state flux cone*. The flux regime is called *bounded*, if it is not unbounded.

For any reaction 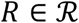, we say *R* is a *blocked reaction* in *G*, if for all flux distributions 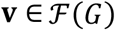, the flux through *R* is zero, namely, *υ_R_* ≡ 0. This is a recurring phenomenon in metabolic networks, in particular when restrictive flux bounds and/or optimization of particular objectives are imposed. As long as we only concern ourselves with steady state analysis, all blocked reactions can be safely removed from the network.

More generally, we say a reaction 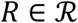 is *fixated* (at some flux value *f*), if for all flux distributions 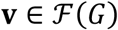, the flux going through *R* is unchanged, namely, 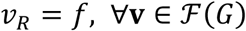. Obviously, the set of blocked reactions is contained in the set of fixated reactions.

### 2.3 Linkage structure and deficiency

Two complexes 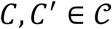 are said to be *directly linked*, if 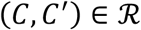 or 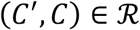. Two complexes 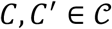 are *linked*, denoted *C* ~ *C’*, if there exist a sequence of complexes (*C* = *C*_*j*_0__, *C*_*j*_1__, …, *C_j_k__* = *C’*), each of which is directly linked to the immediate preceding and succeeding elements. The equivalence relation ~ partitions 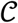 into a family of 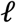 equivalence classes 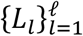, called the *linkage classes* of the network.

For any two complexes 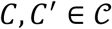, we say *C directly converts to C’*, denoted *C* → *C’*, if 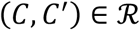 or 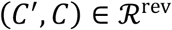. We say *C converts to C’*, denoted *C* ⇒ *C’*, if there exist a sequence of complexes (*C* = *C*_*j*_0__ → *C*_*j*_1__ → … → *C_j_k__* = *C’*). *C* and *C’* are *strongly linked*, denoted *C* ≈ *C’*, if *C* ⇒ *C’* and *C’* ⇒ *C*. The equivalence relation ≈ partitions 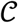 into a family of 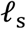 equivalence classes 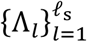, called the *strong linkage classes* of the network.

A *terminal strong linkage class* is a strong linkage class Λ, no complex of which converts to any complex in 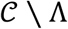. We denote the number of terminal strong linkage classes by 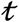. It is trivial to show that, in general, 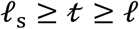. A reaction network is said to be *weakly reversible*, If *C* ~ *C’* implies *C* ≈ *C’*, that is, any two linked complexes convert to each other. In graph-theoretic terms, it means each component of the CRN graph - defined by complexes as vertices and reactions as directed edges – is strongly connected. For weakly reversible networks, every linkage class is a terminal strong linkage class, hence 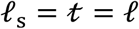.

The *deficiency* of a reaction network is the nonnegative integer, denoted *δ*, defined by (Feinberg, 1987)

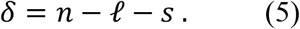

Alternatively, it can be defined as (Gunawardena, 2003; Johnston, 2014)

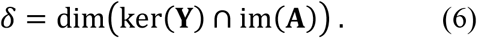

As Eq. (6) demonstrates, the deficiency of a network has to do with the image of stoichiometric flux modes under linear mapping **A**. In this light, for any network *G*, we define the *deficiency space* 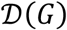 as the set of all feasible complex formation rate vectors, given as follows

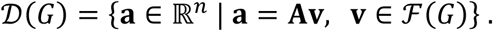

One may interpret the dimension of the deficiency space as a measure of the capacity of a network to permit transformations on the complex level, which are unobserved on the species level, hence may also occur at steady state. This is why the notion of deficiency stands at the center of several seminal results in CRNT, which determine whether special classes of networks may exhibit multistationarity and/or exotic dynamical behavior (Feinberg, 1988).

Given the definition of **A**, it is not surprising that the linkage structure of the network is closely interlinked with the left nullspace of **A**. Let *G* be a closed network comprising 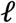 linkage classes 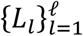. An obvious choice of a basis for the left nullspace of *A*, namely ker **A**^T^, is the set 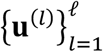, where 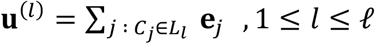. Therefore, the column span of the *complex – linkage class incidence matrix*

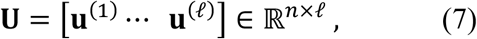

coincides with the left nullspace of **A**. In a similar fashion, we can define the *complex – strong linkage class incidence matrix* as follows

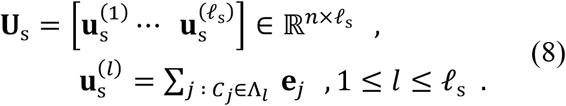

Since every linkage class is a disjoint union of some strong linkage classes, it follows that for any arbitrary network, im(**U**) ⊆ im(**U**_s_), with equality holding only for weakly reversible networks.

## 3 Mathematics of balanced complexes

Let us next give a formal definition of a balanced complex. For a given matrix **X**, let **X**^:*i*^ and **X**^*j*:^ denote the *i*^th^ column and *j*^th^ row of *X*, respectively. A complex 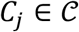 is referred to as a *balanced complex* (BC) for network *G*, if **A**^*j*:^**v** ≡ 0 for all 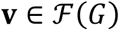. Given the properties of the incidence matrix **A**, this definition complies with the notion that the algebraic sum of fluxes entering complex *C_j_* must be equal to the sum of fluxes leaving it for all feasible distributions 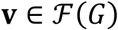, which means this complex has a zero complex formation rate at all steady states.

### 3.1 Factorizations of balanced complexes

As was shown in (Langary, et al., 2021), a complex 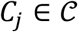 is a BC, if and only if

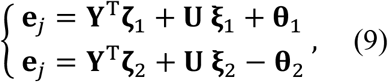

where variables 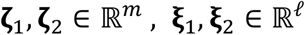 are free parameters, while the parameters 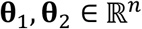 satisfy

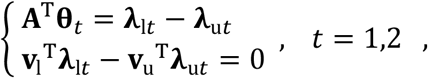

for some nonnegative dual variables 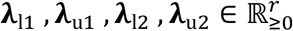 (Langary, et al., 2021).

The set of all balanced complexes in *G*, denoted 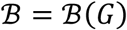, can then be defined as

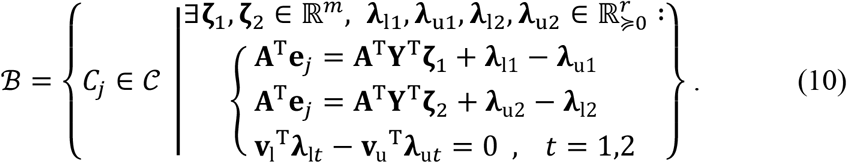

A significant subset of these balanced complexes can be characterized as follows

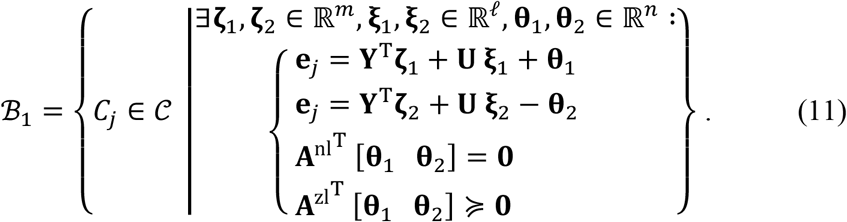

In a canonical flux regime, the above two sets coincide (Langary, et al., 2021), i.e. 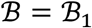. Note that in that case, **A**^nl^ = **A**^rev^ and **A**^zl^ = **A**^irr^.

Furthermore, in a canonical flux regime, if the network is void of any blocked reactions, the set of balanced complexes reduces to

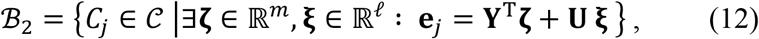

a particular subset of which is given as follows

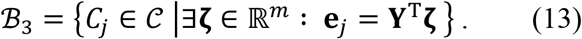

The equalities in Eqs. (10-13), referred to as *factorizations*, explain the formation of a balanced complex 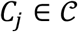 as a combined effect of a number of underlying factors, namely, the stoichiometry (**Y**), linkage structure (**U**), irreversibility patters 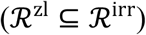 and flux bounds (**v**_l_, **v**_u_).

### 3.2 Nonstoichiometric balanced complexes

It is worth noting that

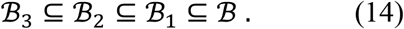

A complex 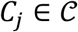 is called a *strictly stoichiometric BC*, if 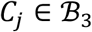; the equality in (13) is referred to as a *strictly stoichiometric factorization* for complex *C_j_*. A complex 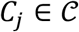 is called a *stoichiometric BC*, if 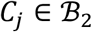; the equality in (12) is a *stoichiometric factorization* for complex *C_j_*.

A 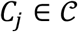 is called a *nonstoichiometric BC*, if 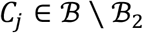; while all nonstoichiometric BCs have an implicit factorization of the form (10), we make a distinction between those which also have an explicit factorization of the form (11), and those which do not. A complex 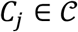 is called a type-I *nonstoichiometric BC*, if 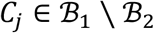, that is, it has a factorization of the form (11), but no stoichiometric factorization. A complex 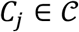 is called a type-II *nonstoichiometric BC*, if 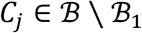, that is, it has a factorization of the form (10), but none of the form (11). A complex 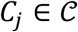 is *unbalanced*, if it has no factorization of the form (10), that is, 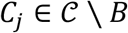.

The emergence of nonstoichiometric BCs is an intriguing phenomenon in (bio)chemical reaction networks. As will become clear in the next section, it reflects on some key structural properties of the reaction network. Furthermore, it also reflects on the reduced capacity of a network to permit variations of the complex formation rate vector at steady state.

## 4 Properties of nonstoichiometric BCs

We hereby aim to seek conditions under which nonstoichiometric BCs may emerge in a network, and to see what implications their existence may have on structural properties of a reaction network. We begin by analyzing type-I nonstoichiometric BCs, which is the only viable type in canonical flux regimes. However, it shall be noted that they may also emerge in non-canonical flux regimes. The following statement was proven in (Langary, et al., 2021).

### Proposition 4.1.

Given a network *G*, if the set 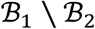 is nonempty, then *G* contains at least two irreversible reactions which are blocked at steady state.

The proof follows from the fact that nonzero (positive) entries in vectors 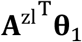 and 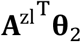 must correspond to blocked reactions in 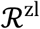. Existence of 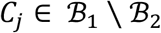 requires that the vectors 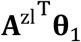 and 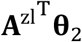 are both nonzero and also not collinear. Proposition 4.1 shows how the presence of a nonstoichiometric BC reflects qualitatively on the steady state properties. In fact, the following statements show that it also contains information about the graphical structure of the CRN.

### Proposition 4.2.

Let the complex 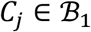 have an explicit factorization of the form (11) with parameters 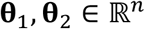. Then, **θ**_1_, **θ**_2_ ∈ im(**U**_s_).

For weakly reversible networks, **U**_s_ = **U**, and hence, im(**U**_s_) = im(**U**). Therefore, the parameter **θ**_t_ can be removed by merging its value with the term **U ξ**_*t*_, which yields a stoichiometric factorization for each 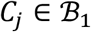. As a result, type-I nonstoichiometric BCs do not emerge in weakly reversible reaction networks.

### Proposition 4.3.

Given a network *G*, if the set 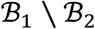 is nonempty, then *G* is not weakly reversible and 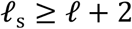.

In particular, it can be shown that blocked irreversible reactions indexed by strictly positive entries of 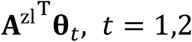 are “bridge reactions” that connect distinct strong linkage classes of the network. In fact, removal of all such blocked reactions increases the number of linkage classes by at least two. Proposition 4.3 shows an interesting contrast to a well-established result in chemical reaction network theory, which states that (full) complex balancing can be obtained at a positive steady state, only if the network is weakly reversible [Proposition 16.5.7 in (Feinberg, 2019)].

Interestingly, once the blocked reactions predicted by Proposition 4.1 are removed, a number of strong linkage classes will be detached from the rest of the network and form separate linkage classes.

### Proposition 4.4.

For a network *G*, let 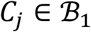. Let *<u>G</u>* be the network obtained by removing all blocked reactions in *G*. Then 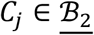, that is, *C_j_* is a stoichiometric BC in *G*. Furthermore, 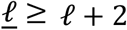.

The following corollary follows immediately from Proposition 4.1.

### Corollary 4.5.

Let *G* be a network with all its blocked reactions have been removed; then we have 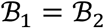 and 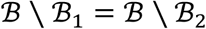; that is, all nonstoichiometric BCs are of type II.

By contrast to type-I nonstoichiometric BCs, the type-II BCs do not necessarily require the network to contain blocked reactions at steady state; hence, they do not automatically follow from modified topological features of the network at steady state. However, a rather similar trend could be observed: For a type-II nonstoichiometric BC to exist, a number of reactions must be fixated at nonzero lower- or upper bounds.

### Proposition 4.6.

Let *G* be a network, all blocked reactions of which have been removed. Suppose 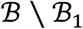 is nonempty. Then *G* contains at least three reactions fixated at a corresponding nonzero lower- or upper bound, for all 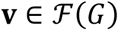.

One can actually come up with a slightly more informative statement.

### Proposition 4.7.

For a network *G*, suppose 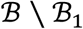 is nonempty. Then *G* contains a reaction 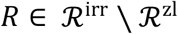 fixated at a positive lower bound. Moreover, there exists another reaction 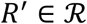 fixated at a positive upper bound, or a reaction 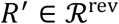 fixated at a negative lower bound.

Note that none of these results imposes any specific restrictions on the graphical structure of the network, unlike the type-I case. For type-II nonstoichiometric BCs, it is not the topological feature but the imposed flux constraints that set the stage for the emergence of additional balanced complexes. The next result follows immediately from Proposition 4.7.

### Corollary 4.8.

Given a network *G*, for the set 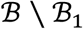 to be nonempty, *G* must operate in a bounded and non-canonical flux regime.

Conversely, suppose *G* is a network operating under an unbounded and/or canonical flux regime, and it contains no blocked reactions. Then, 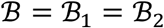.

## 5 Nonstoichiometric BCs and the integration of phantom species

Balanced complexes have been shown to play a key role in the reduction of large-scale metabolic networks [see (Küken, et al., 2021) and references therein]. They shall be viewed as intrinsic properties of the network that shed light on its steady state characteristics regardless of the underlying kinetics governing the conversion of species. The nonstoichiometric BCs have an additional interesting facet to themselves: They carry extra information about the network structure, which is not contained in the steady state equation (3) or in the *linkage closedness condition* **U**^T^**Av** = **0**, but emerges as a joint product of the stoichiometry, graphical structure, and flux constraints.

In what follows, we introduce transformations on a given reaction network *G*, which encodes these extra pieces of information into the steady state equation of a modified network *G*\*, which has the same graphical but a different algebraic structure. This will enable us to exploit *G*\* to make observations about steady-state characteristics of the original CRN *G*. Even though only nonstoichiometric BCs carry extra pieces of information to encode, we shall next begin by introducing these transformations in the more general case, i.e. for any arbitrary BC 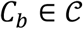.

### 5.1 Insertion of phantom species

Let *G* be a CRN and 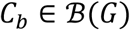 be any balanced complex in *G*, i.e. **e**_*b*_^T^**Av** = 0, 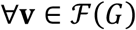. Suppose we ‘amend’ *G* by inserting one molecule of some phantom species *σ* into complex *C_b_*. This yields the modified network 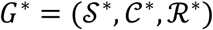, where 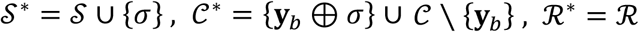. Accordingly, the stoichiometric map and incidence matrix for *G*\* are given by

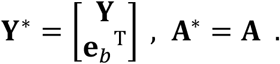

We refer to this transformation as *injection of vector* **e**_*b*_ *into network G*. As this procedure is only applied to some balanced complex *C_b_* and it involves insertion of some new species (not already in the network), the following result is trivially obtained.

#### Lemma 5.1.

Let *G* be a network, 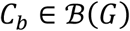 and *G*\* the modified network obtained by injection of **e**_*b*_ into *G*. Then *G* and *G*\* have identical steady state flux distributions, that is, 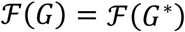. In particular, any balanced complex of one is a balanced complex of the other, 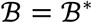. Moreover, we have the inclusion 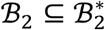, which turns to a strict inclusion, if 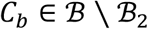.

While *G* and *G*\* have identical balanced complexes, the stoichiometric nature of BCs may differ across the two networks. In particular, for the modified complex, we have 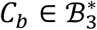, regardless of its original categorization in *G*. For any chosen 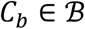, this transformation preserves the steady state flux space, and the information carried by any such BC is hereby encoded into the steady state equation. Clearly, the number of nonstoichiometric BCs strictly decreases in this process, in case of 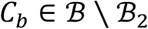.

### 5.2 Characteristics of the modified network

The next question to address is how the stoichiometric subspace develops under the introduced transformation.

#### Proposition 5.2.

Let *G* be a network, 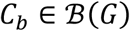 and *G*\* the modified network obtained by injection of **e**_*b*_ into *G*. Let us denote the stoichiometry matrices of *G* and *G*\* by **N** and **N**\*, respectively. Then

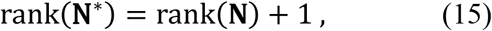

if and only if 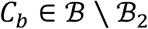. Moreover, rank(**N**\*) = rank(**N**), if and only if 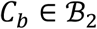.

It is only the injection of nonstoichiometric BCs that increases the rank of the stoichiometric subspace. This yields the following corollary.

#### Corollary 5.3.

Let *G, G*\* be as defined in Proposition 5.2, and 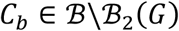. Then, dim ker(**N**\*) = dimker(**N**) – 1.

On the face of it, this seems to contradict the statement of Lemma 5.1. However, the ambiguity is easily sorted out, because not every vector in ker(**N**) lies in 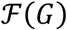, simply due to irreversibility patterns and flux bounds. The apparent difference is only due to the fact that the extra piece of information carried by nonstoichiometric BC *C_b_* is now encoded in the steady state equation **N**\***v** = **0**.

The following result follows readily from the definition of deficiency:

#### Proposition 5.4.

Let *G* be a network, 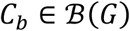 and *G*\* the modified network obtained by injection of **e**_*b*_ into *G*. Let us denote the deficiency of *G* and *G*\* by *δ* and *δ*\*, respectively. Then

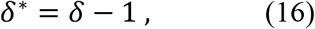

if and only if 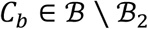. Moreover, *δ*\* = *δ*, if and only if 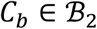.

The deficiency is often characterized as a measure of how tightly the complex formation rate vector, i.e. the image of the steady state flux space under mapping **A**, is constrained at steady state (Feinberg, 2019), that is, the algebraic dimension of the deficiency space. Let us next focus on cases where *C_b_* is a nonstoichiometric BC. Given that *G* and *G*\* have the exact same flux distributions and the exact same incidence matrix, the contrast in their deficiency values is intriguing.

It is worth noting that the deficiency, as defined in (5) or (6), completely neglects of the fine details of the directionality of reactions, or the active bounds on flux through reactions. *The reaction arrows exert their influence only to the extent that they serve to partition the complexes into linkage classes* (Feinberg, 2019). For the rather abstract notion of unconstrained networks, it is trivial to show that the deficiency value coincides with the dimension of the deficiency space, that is

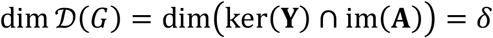

However, in more practical settings, e.g. the constrained framework presented in flux balance analysis of metabolic networks, the deficiency –as defined– should be understood only as an upper bound on the dimension of the deficiency space, that is, the following more general relation holds

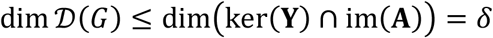

In the case of *G* and *G*\*, the two networks form identical complex formation rate vectors at steady state, that is, have identical deficiency spaces. However, the nominal deficiency of *G, δ*, is larger than that of *G*\*, i.e. *δ*\* = *δ* – 1. This suggests that, due to abovementioned factors, the network *G* is in fact more constrained at steady state than is reflected by *δ* = ker(**Y**) ⋂ im(**A**). Those factors cause the deficiency of *G* to be *effectively* less than *δ*, and no larger than *δ*\*.

## 6 Effective deficiency

Let *G* be a network and 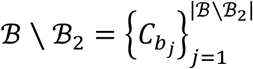 be the set of nonstoichiometric BCs for *G*. Let *G*_1_ be the network obtained by injection of vector **e**_*b*_1__ into *G*. Given the discussion in Section 5, *δ*_1_ = *δ* – 1. Moreover, the equality 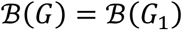 and strict inclusion 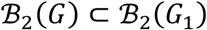 hold. Even though the remaining elements in 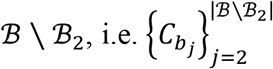 are balanced complexes of *G*_1_, not all of them are necessarily a nonstoichiometric BC for *G*_1_. However, any nonstoichiometric BC of *G*_1_ must lie in 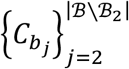.

Now suppose some member of 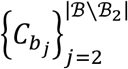 is a nonstoichiometric BC for *G*_1_. Without loss of generality, let it be labelled *C*_*b*_2__. Let *G*_2_ be the network obtained by injection of vector **e**_*b*_2__ into *G*_1_. It follows that *δ*_2_ = *δ*_1_ – 1. We then proceed further by searching the set 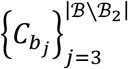 for any complex that may lie in 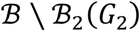. If it exists, we construct *G*_3_ by injection of the corresponding vector, and so on.

As long as the most recent network in the sequence *G* → *G*_1_ → *G*_2_ → … has any nonstoichiometric BC, one can basically repeat this process of inserting distinct phantom species and obtain the next modified network in the sequence. After repeating this process consecutively *d* times, for some d, we will obtain 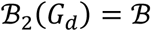; that is, there remains no nonstoichiometric BC left to carry on with the process.

Our first observation is that, by virtue of Lemma 5.1, all networks *G*, *G*_1_, …, *G_d_* have identical steady state flux distributions. Since they all have the same incidence matrix, they also share the same deficiency space, i.e. the steady state variations of the complex formation rate vector is also identical across all such networks. However, by virtue of Proposition 5.4, *δ_d_* = *δ_d-1_* – 1 = *δ_d-2_* – 2 = … = *δ* – *d*. It follows that, due to constraints on the flux space imposed by irreversibility patterns and flux bounds, the deficiency of network *G* is effectively no larger than *δ* – *d*.

In order to be able to define the notion of *effective deficiency* unambiguously, the question of uniqueness needs to be addressed: Had one chosen a different order for the sequence of nonstoichiometric BCs to inject and construct *G* → *G*_1_ → *G*_2_ → …, would one have still obtained the same value for d? The next statement addresses this question.

### Theorem 6.1.

The maximum length of the sequence *G* → *G*_1_ → … → *G_d_*, that is, *d* is independent of the choice and order of nonstoichiometric BCs used to construct it.

Theorem 6.1 paves the way for the definition of effective deficiency for networks under irreversibility constraints and flux bounds.

### Definition

Let *G* be a network and 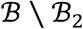 the set of nonstoichiometric BCs for *G*. Let *G*_1_, …, *G_d_* be any arbitrary sequence of modified networks constructed iteratively by means of injecting nonstoichiometric BCs, and suppose 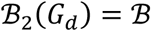. The effective deficiency of *G*, denoted *δ*^eff^, is defined as follows

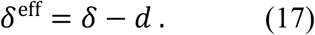

Note that 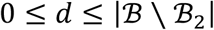. It follows that 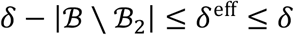. The following result facilitates the computation of effective deficiency, without having to explicitly construct any sequence of modified networks.

### Theorem 6.2.

Suppose 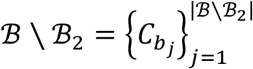 is the set of nonstoichiometric BCs for *G*. Let us construct the matrix 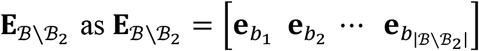. The effective deficiency of *G* is equal to

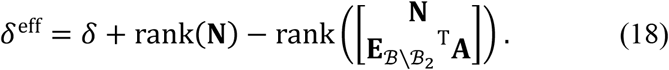

One can seamlessly replace the set 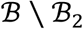 in Proposition 6.2 with the set of all BCs 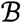 (and replace 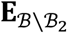 by 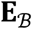, which contains all balancing vectors as columns). Nevertheless, the exact same equality will still hold:

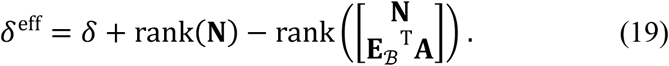

However, the same value of *δ*^eff^ would be obtained at the expense of a higher computational effort. This is due to the fact that injection of stoichiometric BCs has neither an impact on the deficiency of the network nor on its flux distributions, simply because it incorporates no additional information into the stoichiometric structure.

In more technical terms, it can be shown that the effective deficiency has to do with the codimension of im(**Y**^T^) + im(**U**), as stated in the following proposition.

### Proposition 6.3.

Suppose 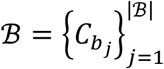 be the set of BCs for *G*, and 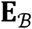 defined as 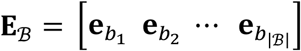. Then, the drop in deficiency, *d*, is the codimension of im(**Y**^T^) + im(**U**) in im(**Y**^T^) + im(**U**) + 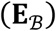, that is

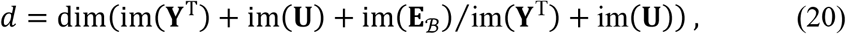

where *V/W* denotes the quotient space of *V* by *W*.

## 7 Some Examples

In Section 6, we introduced the notion of effective deficiency, and established its relation to the existence of nonstoichiometric BCs in the network. In the following, we illustrate this phenomenon via a number of examples. Here, we present toy networks, the design of which does not necessarily reflect real-world chemical or metabolic pathways, but is actually tailored for the purpose of illustration. On that account, the unrealistic nature of these toy networks shall not be viewed critically, as the goal is to simply show a couple of nonstoichiometric BCs and their consequences within a small and representable setting.

As far as real-world applications are concerned, the presence of nonstoichiometric BCs across a wide range of metabolic models has already been confirmed in a recent study (Langary, et al., 2021), which implies the exact same consequences for the effective deficiency of those metabolic networks.

### 7.1 Toy network I: a type-I nonstoichiometric BC

Let us consider the toy network in Fig. 1.

**Fig. 1.**
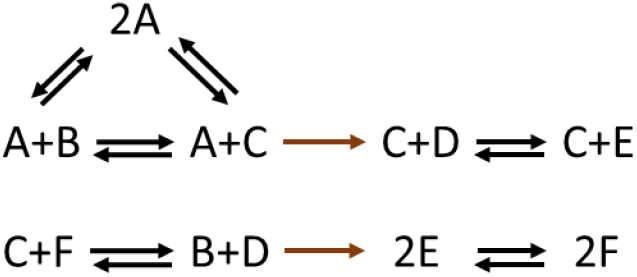
A type-I nonstoichiometric BC in a toy network. The basic conversion diagram depicts a network with *m* = 6 species, *n* = 9 complexes, and *r* = 8 reactions. The network does not contain any stoichiometric BC. However, it contains exactly one type-I nonstoichiometric BC (complex *2A*). In line with the prediction of Proposition 4.1, the network also contains two blocked irreversible reactions (highlighted in brown).

The basic conversion diagram presented in Fig. 1 (including the reactions highlighted in brown) portrays a network operating in a canonical flux regime with *n* = 9 complexes,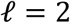 linkage classes and of rank *s* = 5. As a result, the network is of deficiency 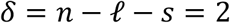. It contains no stoichiometric BCs. However, it can be shown that the complex (*2A*) has a nonstoichiometric factorization of the form Eq. (11), hence, it is a type-I nonstoichiometric BC. The detailed parameter values for the factorization are presented in the Supplementary.

As predicted by Proposition 4.1, the presence of a type-I nonstoichiometric BC is accompanied by having two irreversible reactions blocked at steady state, both of which are highlighted in brown. Moreover, (*2A*) is the only balanced complex, hence, *d* is bound from below and above to be exactly one. Therefore, while the network’s nominal deficiency is two, it follows from the above results that the network is of effective deficiency *δ*^eff^ = *δ* – *d* = 1.

Let us next consider the reduced network obtained by removing all blocked reactions, but including all reactions shown in black. The reduced network will contain no nonstoichiometric BCs, since the balanced complex (*2A*) has turned into a stoichiometric BC in this network. It has *n’* = 9 complexes, 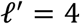 linkage classes and rank *s’* = 4. It follows that the reduced network is of deficiency 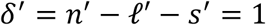. This is in accord with the value calculated for the effective deficiency of the original network.

### 7.2 Toy network II: a type-II nonstoichiometric BC

Next, let us consider the conversion diagram shown in Fig. 2.

**Fig. 2.**
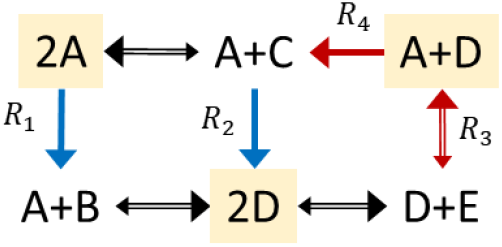
Type-II nonstoichiometric BCs in a toy network. The conversion diagram portrays a network with *m* = 5 species, *n* = 6 complexes, and *r* = 7 reactions. The network operates in a non-canonical and bounded flux regime, where the fluxes of irreversible reactions *R*_1_ and *R*_2_ have strictly positive lower bounds (*υ*_l, 1_ = *υ*_l, 2_ = 100), while the fluxes of reactions *R*_3_ and *R*_4_ have finite upper bounds (*υ*_u, 3_ = *υ*_u, 4_ = 200). The network contains three type-II nonstoichiometric BCs, highlighted in yellow. The rest of complexes are stoichiometric BCs.

For the reversible reactions, the larger arrow size depict the direction of the flux associated with a positive sign. In line with the prediction of Propositions 4.6 and 4.7, the network contains a number of reactions fixated at corresponding lower- and upper bounds, shown in blue and red, respectively.

The toy example in Fig. 2 portrays a network operating in a non-canonical and bounded flux regime. The three complexes (*A* + *B*), (*A* + *C*) and (*D* + *E*) are strictly stoichiometric BCs, due to the fact that species *B, C*, and *E* do not appear elsewhere in the toy network. The network has *n* = 6 complexes, 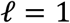 linkage class and rank *s* = 4. Hence, the nominal deficiency of this network is 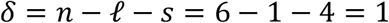.

The other three complexes, that is, (*2A*), (*A* + *D*) and (*2D*) are type-II nonstoichiometric BCs. One can basically associate each with a nonstoichiometric factorization of the form (10) (details are available in the Supplementary). The existence of a nonstoichiometric BC implies that the effective deficiency *δ*^eff^ of this network must be strictly smaller than its nominal deficiency. Coupled with the knowledge that the deficiency only takes nonnegative values, it follows that *δ*^eff^ = 0. In other words, this is effectively a deficiency zero network, and thereby it is complex-balanced.

In line with the statements of Propositions 4.6 and 4.7, the network contains a number of reactions fixated at their lower- and upper bounds, shown in blue and red, respectively. It is worth noting that the formation of nonstoichiometric BCs in this network rely on the irreversibility of *R*_1_ and *R*_2_, the non-canonical flux regime imposed by strictly positive lower bounds on fluxes of of *R*_1_ and *R*_2_, as well as the unbounded flux regime imposed by finite upper bounds on fluxes of of *R*_3_ and *R*_4_. These *imposed constraints* force the network to perform like a deficiency-zero network and hence be complex-balanced, despite the fact that its underlying structure had the capacity to perform like a network of higher deficiency.

### 7.3 Effective deficiency of large-scale metabolic networks

We use twelve genome-scale metabolic networks (Andersen, et al., 2008; Zhang, et al., 2009; Nogales, et al., 2008; Gonzalez, et al., 2010; Quek & Nielsen, 2008; Benedict, et al., 2012; Orth, et al., 2011; Imam, et al., 2015; Feist, et al., 2006; Fang, et al., 2010) (Arnold & Nikoloski, 2014; Lu, et al., 2019) from all kingdoms of life obtained from Küken et al. (2021) to investigate deficiency in networks with two sets of constraints (i) imposing reaction irreversibility constraints as specified in the original model reconstruction, and (ii) imposed optimality of specific growth rate in addition to reaction reversibility constraints. Note that blocked reactions are removed from the networks before BC detection and, therefore, the networks do not contain type-I nonstoichiometric BCs. In other words, all type-I nonstoichiometric BCs are transformed into stoichiometric BCs as a result of these removals (see Proposition 4.4). In this setting, we compare the structural deficiency and effective deficiencies obtained under the two different sets of constraints. The structural deficiency ranges from 57 for *T. maritima* to 1097 for *S. cerevisiae* (Fig. 3). We find that the effective deficiency obtained from scenario (i) is the same with the structural deficiency for the networks of *A. thaliana, M. musculus, N. pharaonis*, and *P. putida.* For the remaining eight networks, we find a reduction of the effective deficiency in comparison to the structural deficiency, ranging from 1.8% in *E. coli* and 35.3% in *M. barkeri* (Fig. 3). Considering the additional constraint on the specific growth rate, fixed at its optimum, in scenario (ii) we observe the effective deficiency to be smaller than the structural deficiency in all networks, with the smallest decrease in networks of *A. thaliana* (0.8%), *E. coli* (1.8%), *M. musculus* (3.6%) and *A. niger* (4.2%) and the largest decrease in the networks of *M. barkeri* (35.3%), *M. acetivorans* (11%), *C. reinhardtii* (9.9%) and *T. maritima* (9.6%).

**Fig. 3.**
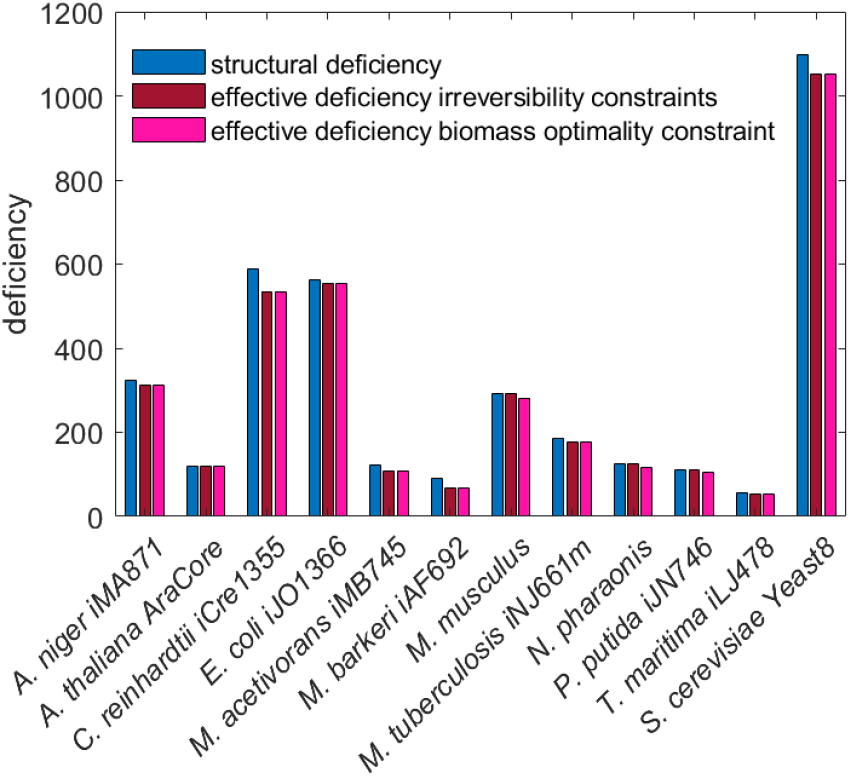
Structural and effective deficiency of real-world metabolic networks. Balanced complexes are identified in networks of twelve species from all kingdoms of life under two different scenarios: (i) imposing reaction irreversibility constraints, and (ii) considering in addition specific growth rate optimality. The effective deficiency is smaller than the structural deficiency when the additional constraint on optimality of specific growth rate is imposed for all networks.

## 8 Conclusion

Here we introduced the notion of effective deficiency that takes into account not only the structure of the network but also its operational constraints in the calculation of deficiency. The effective deficiency relies on the presence of nonstoichiometric balanced complexes, which we have shown to be present in large-scale metabolic networks across kingdoms of life. In addition, our results point at a subtle relation between effective deficiency and robustness of some reaction fluxes. Future work will aim at employing these findings in characterizing classes of networks that exhibit particular dynamical properties that can be ensured or precluded based on the notion of effective deficiency.

## Declarations

### Availability of data and material

(All networks used in the analysis are available here)

### Code availability

(The code is available here)

### Authors’ contributions

(Conceptualization: Z.N., D.L.; Methodology: Z.N., D.L.; Formal analysis and investigation: D.L.; Implementation: A.K.; Writing - original draft preparation: D.L.; Writing - review and editing: D.L., A.K., Z.N.; Funding acquisition: Z.N.; Resources: A.K.; Supervision: Z.N.)

## Supplementary Information

Proofs for the theorems and propositions in the text, as well as detailed nonstoichiometric factorizations for the toy examples in Section 7 are provided in <u>Supplementary Information [link]</u>.

