## Supplementary Information for "Nonstoichiometric balanced complexes: Implications on the effective deficiency of the underlying metabolic network"

### Supplementary Information
### for
### Nonstoichiometric balanced complexes:
### Implications on the effective deficiency of the underlying metabolic network

Damoun Langary[1,2], Anika Küken[2], Zoran Nikoloski[1,2]

[1]Systems Biology and Mathematical Modeling, Max Planck Institute of Molecular Plant Physiology, Potsdam, Germany

[2]Bioinformatics, Institute of Biochemistry and Biology, University of Potsdam, Potsdam, Germany

#### S1 Proofs of statements

**Proposition 4.1**. For a network $G$, let the set $\mathcal{B}_1 \setminus \mathcal{B}_2$ be nonempty. Then, $G$ contains at least two irreversible reactions which are blocked at steady state.

A proof for Proposition 4.1 is given in (Langary, et al., 2021).■

To facilitate proving the next statement, let us first present the following lemma.

**Lemma S1.** Let the complex $C_b \in \mathcal{B}_1$ have an explicit factorization of the form (11) with parameters $\boldsymbol{\theta}_1, \boldsymbol{\theta}_2 \in \mathbb{R}^n$. Let $C_i$ and $C_j$ be arbitrary complexes in the network, which satisfy $C_i \Rightarrow C_j$. Then, $\theta_{t,i} \le \theta_{t,j}\ \ ,\ t = 1,2$.

<u>Proof</u>: Given $C_i \Rightarrow C_j$, there exists a path $C_i = C_{j_0} \to C_{j_1} \to \cdots \to C_{j_\kappa} = C_j$ . Now, given Eq. (11), for any irreversible reaction with zero lower bound $R_k :\ C_{k_\mathrm{r}} \rightharpoonup C_{k_\mathrm{p}} \in \mathcal{R}^{\mathrm{zl}} \subseteq \mathcal{R}^{\mathrm{irr}}$, we obtain $\mathbf{a}_k{}^{\mathrm{T}}\boldsymbol{\theta}_t \ge 0$ ; hence,

$$\theta_{t,k_\mathrm{r}} \le \theta_{t,k_\mathrm{p}} .$$

For any other reaction (reversible, or irreversible with nonzero lower bound, $\mathbf{a}_k{}^{\mathrm{T}}\boldsymbol{\theta}_t = 0$ ; hence,

$$\theta_{t,k_\mathrm{r}} = \theta_{t,k_\mathrm{p}} .$$

Any direct conversion $C_{j_\alpha} \to C_{j_{\alpha+1}}$ in the chain $C_i = C_{j_0} \to C_{j_1} \to \cdots \to C_{j_\kappa} = C_j$ corresponds to either a reversible or an irreversible reaction. Hence, either $\theta_{t,j_\alpha} \le \theta_{t,j_{\alpha+1}}$ or $\theta_{t,j_\alpha} = \theta_{t,j_{\alpha+1}}$. It follows that

$$\theta_{t,i} \le \theta_{t,j} . \blacksquare$$

**Proposition 4.2**. Let the complex $C_b \in \mathcal{B}_1$ have an explicit factorization of the form (11) with parameters $\boldsymbol{\theta}_1, \boldsymbol{\theta}_2 \in \mathbb{R}^n$. Then, $\boldsymbol{\theta}_1, \boldsymbol{\theta}_2 \in \mathrm{im}(\mathbf{U}_\mathrm{s})$.

Proof: Let $C_i$ and $C_j$ be two complexes in the same strong linkage class. It follows from the definition that $C_i \Rightarrow C_j$ and $C_j \Rightarrow C_i$. As a result, by virtue of Lemma S1, we must have $\theta_{t,i} \leq \theta_{t,j}$ and $\theta_{t,i} \leq \theta_{t,j}$ for $t = 1,2$, which yields $\theta_{t,i} = \theta_{t,j}$. Hence, for all complexes in a single strong linkage class, their corresponding entries in $\boldsymbol{\theta}_t$ have equal values. Therefore, $\boldsymbol{\theta}_t$ can be written as

$$\boldsymbol{\theta}_t = \sum_{l=1}^{\ell_s} \eta_{t,l}\, \mathbf{u}_{\mathrm{s}}^{(l)} = \mathbf{U}_{\mathrm{s}} \boldsymbol{\eta}_t \in \mathrm{im}(\mathbf{U}_{\mathrm{s}}), \qquad t = 1,2\,. \blacksquare \qquad \text{(S1)}$$

**Proposition 4.3**. For a network $G$, let the set $\mathcal{B}_1 \setminus \mathcal{B}_2$ be nonempty. Then, $G$ is not weakly reversible. Moreover, $\ell_{\mathrm{s}} \geq \ell + 2$ .

Proof: [By contradiction] Suppose $G$ is weakly reversible, and suppose $C_b$ be any arbitrary BC in $\mathcal{B}_1$. $C_b$ has a factorization of form (11); furthermore, by virtue of Proposition 4.2, $\boldsymbol{\theta}_t \in \mathrm{im}(\mathbf{U}_{\mathrm{s}})$, $t = 1,2$ . Since $G$ is assumed weakly reversible, every linkage class is a strong linkage class; hence, $\mathbf{U}_{\mathrm{s}} = \mathbf{U}$. As a result,

$$\exists \boldsymbol{\eta}_1 \colon \boldsymbol{\theta}_1 = \mathbf{U}\, \boldsymbol{\eta}_1\,.$$

Therefore,

$$\mathbf{e}_b = \mathbf{Y}^{\mathrm{T}} \boldsymbol{\zeta}_1 + \mathbf{U}\, \boldsymbol{\xi}_1 + \boldsymbol{\theta}_1 = \mathbf{Y}^{\mathrm{T}} \boldsymbol{\zeta}_1 + \mathbf{U}\, (\boldsymbol{\xi}_1 + \boldsymbol{\eta}_1)$$

Hence, $C_b$ has a factorization of the form (12), which means $C_b \in \mathcal{B}_2$, by definition. As a result, $\mathcal{B}_1 \setminus \mathcal{B}_2$ is empty, which is a contradiction.

Consequently, if $\mathcal{B}_1 \setminus \mathcal{B}_2$ is nonempty, then $G$ cannot be weakly reversible; therefore, $\ell_{\mathrm{s}} \geq \ell + 1$. Now, either we have $\ell_{\mathrm{s}} = \ell + 1$ or $\ell_{\mathrm{s}} \geq \ell + 2$. We will next show that the case $\ell_{\mathrm{s}} = \ell + 1$ leads to a contradiction.

Now, suppose $C_b \in \mathcal{B}_1 \setminus \mathcal{B}_2$ and $\ell_s = \ell + 1$. This means every linkage class is a strong linkage class, except for a single linkage class, which consists of two strong linkage classes. From Eq. (11), we have

$$\begin{cases} \mathbf{e}_b = \mathbf{Y}^{\mathrm{T}} \boldsymbol{\zeta}_1 + \mathbf{U}\, \boldsymbol{\xi}_1 + \boldsymbol{\theta}_1 \\ \mathbf{e}_b = \mathbf{Y}^{\mathrm{T}} \boldsymbol{\zeta}_2 + \mathbf{U}\, \boldsymbol{\xi}_2 - \boldsymbol{\theta}_2 \end{cases}. \qquad \text{(S2)}$$

From these factorizations, we can calculate $\mathbf{A}^{\mathrm{T}} \mathbf{e}_b$ as

$$\begin{cases} \mathbf{A}^{\mathrm{T}} \mathbf{e}_b = \mathbf{A}^{\mathrm{T}} \mathbf{Y}^{\mathrm{T}} \boldsymbol{\zeta}_1 + \mathbf{A}^{\mathrm{T}} \boldsymbol{\theta}_1 \\ \mathbf{A}^{\mathrm{T}} \mathbf{e}_b = \mathbf{A}^{\mathrm{T}} \mathbf{Y}^{\mathrm{T}} \boldsymbol{\zeta}_2 - \mathbf{A}^{\mathrm{T}} \boldsymbol{\theta}_2 \end{cases}. \qquad \text{(S3)}$$

Given the other constraints in Eq. (11), we know $\mathbf{A}^{\mathrm{T}} \boldsymbol{\theta}_{\mathrm{t}}$ can only have nonzero values for "bridge reactions" that connect two strong linkage classes within a single linkage class. Moreover, given that $\boldsymbol{\theta}_t \in \mathrm{im}(\mathbf{U}_{\mathrm{s}})$, the vector $\mathbf{A}^{\mathrm{T}} \boldsymbol{\theta}_{\mathrm{t}}$ will have the exact same entry value for all these bridge reactions. It follows that $\mathbf{A}^{\mathrm{T}}\boldsymbol{\theta}_1$ and $\mathbf{A}^{\mathrm{T}}\boldsymbol{\theta}_2$ are collinear vectors, with a positive ratio.[i]

Let $\mathbf{A}^{\mathrm{T}}\boldsymbol{\theta}_2 = \beta\,\mathbf{A}^{\mathrm{T}}\boldsymbol{\theta}_1$. It follows from Eq. (S3) that

$$(1+\beta)\mathbf{A}^{\mathrm{T}}\boldsymbol{\theta}_1 = \mathbf{A}^{\mathrm{T}}\mathbf{Y}^{\mathrm{T}}(\boldsymbol{\zeta}_2 - \boldsymbol{\zeta}_1)\,. \qquad \text{(S4)}$$

Since $\beta$ is positive, it follows that

$$\mathbf{A}^{\mathrm{T}}\boldsymbol{\theta}_1 = \frac{1}{1+\beta}\mathbf{A}^{\mathrm{T}}\mathbf{Y}^{\mathrm{T}}(\boldsymbol{\zeta}_2 - \boldsymbol{\zeta}_1)\,.$$

Hence, there exist a vector $\boldsymbol{\xi}'$ such that,

$$\boldsymbol{\theta}_1 = \frac{1}{1+\beta}\mathbf{Y}^{\mathrm{T}}(\boldsymbol{\zeta}_2 - \boldsymbol{\zeta}_1) + \mathbf{U}\,\boldsymbol{\xi}'. \qquad \text{(S5)}$$

Replacing Eq. (S5) in Eq. (S1) will yield a stoichiometric factorization for $C_b$, which is a contradiction. Therefore, $\ell_{\mathrm{s}} \geq \ell + 2$. ■

**Proposition 4.4**. For a network $G$, let $C_b \in \mathcal{B}_1$. Let $\underline{G}$ be the network obtained by removing all blocked reactions in $G$. Then $C_b \in \underline{\mathcal{B}_2}$, that is, $C_b$ is a stoichiometric BC in $\underline{G}$. Furthermore, $\underline{\ell} \geq \ell + 2$ .

<u>Proof</u>: It was shown in proof of Proposition 4.1 that all reactions corresponding to nonzero (positive) entries of $\mathbf{A}^{\mathrm{T}}\boldsymbol{\theta}_{\mathrm{t}}, t = 1{,}2$ are blocked at steady state. It follows that once all blocked reactions are removed, we obtain $\mathbf{a}_k{}^{\mathrm{T}}\boldsymbol{\theta}_t = 0$ for all the remaining reactions. On the other hand, $\mathrm{im}(\underline{\mathbf{A}}) \subset \mathrm{im}(\mathbf{A})$; hence, $\ker(\mathbf{A}^{\mathrm{T}}) \subset \ker(\underline{\mathbf{A}}^{\mathrm{T}})$, that is, $\mathrm{im}(\mathbf{U}) \subset \mathrm{im}(\underline{\mathbf{U}})$. It follows that Eq. (S2) can be written as

$$\begin{cases} \mathbf{e}_b = \mathbf{Y}^{\mathrm{T}}\boldsymbol{\zeta}_1 + \underline{\mathbf{U}}\,\boldsymbol{\xi}_1' + \boldsymbol{\theta}_1 \\ \mathbf{e}_b = \mathbf{Y}^{\mathrm{T}}\boldsymbol{\zeta}_2 + \underline{\mathbf{U}}\,\boldsymbol{\xi}_2' - \boldsymbol{\theta}_2 \end{cases}. \qquad \text{(S6)}$$

Multiplying both equations by $\underline{\mathbf{A}}^{\mathrm{T}}$ from the left, we obtain

$$\begin{cases} \underline{\mathbf{A}}^{\mathrm{T}}\mathbf{e}_b = \underline{\mathbf{A}}^{\mathrm{T}}\mathbf{Y}^{\mathrm{T}}\boldsymbol{\zeta}_1 + \underline{\mathbf{A}}^{\mathrm{T}}\boldsymbol{\theta}_1 \\ \underline{\mathbf{A}}^{\mathrm{T}}\mathbf{e}_b = \underline{\mathbf{A}}^{\mathrm{T}}\mathbf{Y}^{\mathrm{T}}\boldsymbol{\zeta}_2 - \underline{\mathbf{A}}^{\mathrm{T}}\boldsymbol{\theta}_2 \end{cases}. \qquad \text{(S7)}$$

However, $\underline{\mathbf{A}}^{\mathrm{T}}\boldsymbol{\theta}_t = \mathbf{0}$ , $t = 1{,}2$ as we explained. Therefore,

$$\begin{cases} \underline{\mathbf{A}}^{\mathrm{T}}\mathbf{e}_b = \underline{\mathbf{A}}^{\mathrm{T}}\mathbf{Y}^{\mathrm{T}}\boldsymbol{\zeta}_1 \\ \underline{\mathbf{A}}^{\mathrm{T}}\mathbf{e}_b = \underline{\mathbf{A}}^{\mathrm{T}}\mathbf{Y}^{\mathrm{T}}\boldsymbol{\zeta}_2 \end{cases}, \qquad \text{(S8)}$$

[i] Neither $\mathbf{A}^{\mathrm{T}}\boldsymbol{\theta}_1$ nor $\mathbf{A}^{\mathrm{T}}\boldsymbol{\theta}_2$ can be zero; because in such cases, obviously, $C_b \in \mathcal{B}_2$.

which implies has a stoichiometric factorization in $\underline{G}$. Furthermore, the fact that $\underline{\ell} \geq \ell + 2$ follows directly from $\ell_{\mathrm{s}} \geq \ell + 2$, and that $\mathbf{A}^{\mathrm{T}}\boldsymbol{\theta}_{\mathrm{t}}\,, t = 1{,}2$ must have nonzero values for two distinct class of bridge reactions, in order to be linearly dependent.■

**Corollary 4.5**. Let $G$ be a network with all its blocked reactions have been removed; then we have $\mathcal{B}_1 = \mathcal{B}_2$ and $\mathcal{B} \setminus \mathcal{B}_1 = \mathcal{B} \setminus \mathcal{B}_2$ ; that is, all nonstoichiometric BCs are of type II.

Proof: It is straightforward to prove this statement by contradiction, using Proposition 4.1.■

**Proposition 4.6**. Let $G$ be a network, all blocked reactions of which have been removed. Suppose $\mathcal{B} \setminus \mathcal{B}_1$ is nonempty. Then $G$ contains at least three reactions fixated at a corresponding nonzero lower- or upper bound, for all $\mathbf{v} \in \mathcal{F}(G)$.

A proof for Proposition 4.6 is given in (Langary, et al., 2021). Given the general factorization of $C_b \in \mathcal{B} \setminus \mathcal{B}_1$ as follows

$$\begin{cases} \mathbf{A}^{\mathrm{T}}\mathbf{e}_b = \mathbf{A}^{\mathrm{T}}\mathbf{Y}^{\mathrm{T}}\boldsymbol{\zeta}_1 + \boldsymbol{\lambda}_{\mathrm{l1}} - \boldsymbol{\lambda}_{\mathrm{u1}} \\ \mathbf{A}^{\mathrm{T}}\mathbf{e}_b = \mathbf{A}^{\mathrm{T}}\mathbf{Y}^{\mathrm{T}}\boldsymbol{\zeta}_2 + \boldsymbol{\lambda}_{\mathrm{u2}} - \boldsymbol{\lambda}_{\mathrm{l2}} \\ \mathbf{v}_{\mathrm{l}}^{\mathrm{T}}\boldsymbol{\lambda}_{\mathrm{l}t} - \mathbf{v}_{\mathrm{u}}^{\mathrm{T}}\boldsymbol{\lambda}_{\mathrm{u}t} = 0\ ,\quad t = 1{,}2 \end{cases}, \qquad (S9)$$

the proof evokes the exact same argument used in proof of Proposition 4.4 to show that existence of a nonstoichiometric BC $C_b \in \mathcal{B} \setminus \mathcal{B}_1$ requires having both vectors $\boldsymbol{\lambda}_{\mathrm{l1}} - \boldsymbol{\lambda}_{\mathrm{u1}}$ and $\boldsymbol{\lambda}_{\mathrm{u2}} - \boldsymbol{\lambda}_{\mathrm{l2}}$ nonzero and linearly independent. The fixated reactions are those associated with nonzero entries in $\boldsymbol{\lambda}_{\mathrm{l1}}, \boldsymbol{\lambda}_{\mathrm{u1}}, \boldsymbol{\lambda}_{\mathrm{u2}}, \boldsymbol{\lambda}_{\mathrm{l2}}$.■

**Proposition 4.7**. For a network $G$, suppose $\mathcal{B} \setminus \mathcal{B}_1$ is nonempty. Then $G$ contains a reaction $R \in \mathcal{R}^{\mathrm{irr}} \setminus \mathcal{R}^{\mathrm{zl}}$ fixated at a positive lower bound. Moreover, there exists another reaction $R' \in \mathcal{R}$ fixated at a positive upper bound, or a reaction $R' \in \mathcal{R}^{\mathrm{rev}}$ fixated at a negative lower bound.

Proof: It has been established that $\boldsymbol{\lambda}_{\mathrm{l1}} - \boldsymbol{\lambda}_{\mathrm{u1}}$ is nonzero, while the variables $\boldsymbol{\lambda}_{\mathrm{l1}},\ \boldsymbol{\lambda}_{\mathrm{u1}}$ are bound to be nonnegative. Next, let us focus on the third equality in (S9):

$$\mathbf{v}_{\mathrm{l}}^{\mathrm{T}}\boldsymbol{\lambda}_{\mathrm{l1}} - \mathbf{v}_{\mathrm{u}}^{\mathrm{T}}\boldsymbol{\lambda}_{\mathrm{u1}} = \sum_{R\in\mathcal{R}^{\mathrm{zl}}} v_{\mathrm{l},R}\lambda_{\mathrm{l1},R} + \sum_{R\in\mathcal{R}^{\mathrm{irr}}\setminus\mathcal{R}^{\mathrm{zl}}} v_{\mathrm{l},R}\lambda_{\mathrm{l1},R} + \sum_{R\in\mathcal{R}^{\mathrm{rev}}} v_{\mathrm{l},R}\lambda_{\mathrm{l1},R} + \sum_{R\in\mathcal{R}} - v_{\mathrm{u},R}\lambda_{\mathrm{u1},R} = 0\,. \qquad (S10)$$

Now, the 1st term $\sum_{R\in\mathcal{R}^{\mathrm{zl}}} v_{\mathrm{l},R}\lambda_{\mathrm{l1},R}$ is by definition of $\mathcal{R}^{\mathrm{zl}}$ equal to zero. The 2nd term $\sum_{R\in\mathcal{R}^{\mathrm{irr}}\setminus\mathcal{R}^{\mathrm{zl}}} v_{\mathrm{l},R}\lambda_{\mathrm{l1},R}$ can only take nonnegative values for $\boldsymbol{\lambda}_{\mathrm{l1}} \geqslant \mathbf{0}$, while the 3rd and 4th terms can only take nonpositive values. This follows from the fact that

$$v_{\mathrm{l},R} > 0, \qquad \forall R \in \mathcal{R}^{\mathrm{irr}} \setminus \mathcal{R}^{\mathrm{zl}}\,;$$
$$v_{\mathrm{l},R} < 0, \qquad \forall R \in \mathcal{R}^{\mathrm{rev}}\,;$$
$$v_{\mathrm{u},R} > 0, \qquad \forall R \in \mathcal{R}\,.$$

For the above sum to equal zero, while having a nonzero vector $\boldsymbol{\lambda}_{\mathrm{l}1} - \boldsymbol{\lambda}_{\mathrm{u}1}$, it is essential to have the strict inequality $\sum_{R\in\mathcal{R}^{\mathrm{irr}}\setminus\mathcal{R}^{\mathrm{zl}}} v_{\mathrm{l},R}\lambda_{\mathrm{l}1,R} > 0$, which in turn implies that

$$\exists R \in \mathcal{R}^{\mathrm{irr}} \setminus \mathcal{R}^{\mathrm{zl}}\,;\;\; \lambda_{\mathrm{l}1,R} > 0\,.$$

This completes the proof for the first statement of Proposition 4.7.

For the second part, note that the equality in (S10) and the fact that $\sum_{R\in\mathcal{R}^{\mathrm{irr}}\setminus\mathcal{R}^{\mathrm{zl}}} v_{\mathrm{l},R}\lambda_{\mathrm{l}1,R} > 0$ implies that

$$\sum_{R\in\mathcal{R}^{\mathrm{rev}}} v_{\mathrm{l},R}\lambda_{\mathrm{l}1,R} < 0 \quad \text{or} \quad \sum_{R\in\mathcal{R}} - v_{\mathrm{u},R}\lambda_{\mathrm{u}1,R} < 0\,.$$

Consequently,

$$\exists R \in \mathcal{R}^{\mathrm{rev}}\,;\;\; \lambda_{\mathrm{l}1,R} > 0 \quad \text{or} \quad \exists R \in \mathcal{R}\,;\;\; \lambda_{\mathrm{u}1,R} > 0\,,$$

which finishes the second part of the proof.■

**Corollary 4.8**. Given a network $G$, for the set $\mathcal{B} \setminus \mathcal{B}_1$ to be nonempty, $G$ must operate in a bounded and non-canonical flux regime.

Conversely, suppose $G$ is a network operating under an unbounded and/or canonical flux regime, and it contains no blocked reactions. Then, $\mathcal{B} = \mathcal{B}_1 = \mathcal{B}_2$.

Proof: This simply follows from Proposition 4.7. A nonempty set $\mathcal{B} \setminus \mathcal{B}_1$ means at least one irreversible reaction is fixated at a positive lower bound, which itself means the network is not operating under a canonical flux regime. The 2nd statement of Proposition 4.7 then implies that the network is not operating in an unbounded flux regime.

Conversely, suppose $G$ is a network operating under an unbounded and/or canonical flux regime, then $\mathcal{B} \setminus \mathcal{B}_1$ must be empty. Given the general inclusion $\mathcal{B}_3 \subseteq \mathcal{B}_2 \subseteq \mathcal{B}_1 \subseteq \mathcal{B}$, this implies $\mathcal{B} = \mathcal{B}_1$. If all blocked reactions are additionally removed, then by virtue of Corollary 4.5, $\mathcal{B}_1 = \mathcal{B}_2$ , which completes the proof.■

**Lemma 5.1** Let $G$ be a network, $C_b \in \mathcal{B}(G)$ and $G^*$ the modified network obtained by injection of $\mathbf{e}_b$ into $G$. Then $G$ and $G^*$ have identical steady state flux distributions, that is, $\mathcal{F}(G) = \mathcal{F}(G^*)$. In particular, any balanced complex of one is a balanced complex of the other, $\mathcal{B} = \mathcal{B}^*$. Moreover, we have the inclusion $\mathcal{B}_2 \subseteq \mathcal{B}_2^*$ turns to a strict inclusion, if $C_b \in \mathcal{B} \setminus \mathcal{B}_2$.

Proof: Let $\mathbf{v}$ be any vector in $\mathcal{F}(G)$. Then, by definition $\mathbf{e}_b{}^{\mathrm{T}}\mathbf{A}\mathbf{v} = 0$. Moreover, the steady state condition implies that $\mathbf{Y}\,\mathbf{A}\,\mathbf{v} = \mathbf{0}$. For the modified network $G^*$,

$$\mathbf{Y}^*\mathbf{A}^*\mathbf{v} = \begin{bmatrix} \mathbf{Y} \\ \mathbf{e}_b{}^{\mathrm{T}} \end{bmatrix} \mathbf{A}\,\mathbf{v} = \begin{bmatrix} \mathbf{Y}\,\mathbf{A}\,\mathbf{v} \\ \mathbf{e}_b{}^{\mathrm{T}}\mathbf{A}\,\mathbf{v} \end{bmatrix} = \mathbf{0} \ .$$

Since $G$ and $G^*$ share the same flux bounds, it follows that $\mathbf{v} \in \mathcal{F}(G^*)$. Therefore, $\mathcal{F}(G) \subseteq \mathcal{F}(G^*)$. Conversely, if $\mathbf{v} \in \mathcal{F}(G^*)$, the steady state equation implies that $\mathbf{v} \in \mathcal{F}(G)$, that is, $\mathcal{F}(G^*) \subseteq \mathcal{F}(G)$. Hence, $\mathcal{F}(G) = \mathcal{F}(G^*)$.

The fact that the two networks $G$ and $G^*$ have identical sets of balanced complexes is an immediate result of $\mathcal{F}(G) = \mathcal{F}(G^*)$. It remains is to show $\mathcal{B}_2 \subseteq \mathcal{B}_2^*$.

Given the fact that the row space of $\mathbf{Y}$ is a subset of the row space of $\mathbf{Y}^*$ and $\mathbf{U} = \mathbf{U}^*$, it is trivial to see any vector in $\mathcal{B}_2$ is a member of $\mathcal{B}_2^*$, hence, $\mathcal{B}_2 \subseteq \mathcal{B}_2^*$. Moreover, if $C_b \in \mathcal{B} \setminus \mathcal{B}_2$, then $\mathcal{B}_2^*$ contains at least one member ($C_b$), which is not in $\mathcal{B}_2$; thereby, we obtain the strict inclusion $\mathcal{B}_2 \subset \mathcal{B}_2^*$.■

**Proposition 5.2**. Let $G$ be a network, $C_b \in \mathcal{B}(G)$ and $G^*$ the modified network obtained by injection of $\mathbf{e}_b$ into $G$. Let us denote the stoichiometry matrices of $G$ and $G^*$ by $\mathbf{N}$ and $\mathbf{N}^*$, respectively. Then

$$\mathrm{rank}(\mathbf{N}^*) = \mathrm{rank}(\mathbf{N}) + 1 \,,$$

if and only if $C_b \in \mathcal{B} \setminus \mathcal{B}_2$. Moreover, $\mathrm{rank}(\mathbf{N}^*) = \mathrm{rank}(\mathbf{N})$, if and only if $C_b \in \mathcal{B}_2$.

Proof: From the definition of $\mathbf{N}^*$

$$\mathbf{N}^* = \mathbf{Y}^*\mathbf{A}^* = \begin{bmatrix} \mathbf{N} \\ \mathbf{e}_b{}^{\mathrm{T}}\mathbf{A} \end{bmatrix} \qquad \text{(S11)} .$$

$\mathbf{N}^*$ contains all the rows of $\mathbf{N}$, plus an additional row $\mathbf{e}_b{}^{\mathrm{T}}\mathbf{A}$. It follows that either $\mathrm{rank}(\mathbf{N}^*) = \mathrm{rank}(\mathbf{N})$ or we have $\mathrm{rank}(\mathbf{N}^*) = \mathrm{rank}(\mathbf{N}) + 1$. As a result, it suffices to prove only one of the above two statements; the other one would follow automatically.

Now suppose $\mathrm{rank}(\mathbf{N}^*) = \mathrm{rank}(\mathbf{N})$. Then, the last row in $\mathbf{N}^*$ must be a linear combination of the rows of $\mathbf{N}$, i.e.

$$\exists \boldsymbol{\mu} : \ \mathbf{e}_b{}^{\mathrm{T}}\mathbf{A} = \boldsymbol{\mu}^{\mathrm{T}}\mathbf{N} = \boldsymbol{\mu}^{\mathrm{T}}\mathbf{Y}\,\mathbf{A} \,,$$

Hence

$$\mathbf{A}^{\mathrm{T}}(\mathbf{e}_b - \mathbf{Y}^{\mathrm{T}}\boldsymbol{\mu}) = \mathbf{0} \,.$$

Given the well-known structure of the nullspace of $\mathbf{A}^{\mathrm{T}}$,

$$\exists \xi_1, \xi_2, \cdots, \xi_\ell \ : \quad \mathbf{e}_b - \mathbf{Y}^{\mathrm{T}}\boldsymbol{\mu} = \sum_{l=1}^{\ell} \xi_l \mathbf{u}_l = \mathbf{U}\,\boldsymbol{\xi}.$$

This means $\mathbf{e}_b$ has a stoichiometric factorization for $G$. Conversely, if $\mathbf{e}_b$ encodes a stoichiometric BC $C_b \in \mathcal{B}_2$ for $G$ ; $\mathbf{e}_b = \mathbf{Y}^{\mathrm{T}}\boldsymbol{\zeta} + \mathbf{U}\,\boldsymbol{\xi}$ ; hence

$$\mathbf{A}^{\mathrm{T}}\mathbf{e}_b = \mathbf{A}^{\mathrm{T}}\mathbf{Y}^{\mathrm{T}}\boldsymbol{\zeta} + \mathbf{A}^{\mathrm{T}}\mathbf{U}\,\boldsymbol{\xi} = \mathbf{A}^{\mathrm{T}}\mathbf{Y}^{\mathrm{T}}\boldsymbol{\zeta} \ .$$

$$\mathbf{e}_b{}^{\mathrm{T}}\mathbf{A} = \boldsymbol{\zeta}^{\mathrm{T}}\mathbf{Y}\,\mathbf{A} = \boldsymbol{\zeta}^{\mathrm{T}}\mathbf{N}\ .$$

Therefore, the last row in $\mathbf{N}^*$ is a linear combination of the rows of **N**, i.e. **N** and $\mathbf{N}^*$ have the exact same row span. Therefore,

$$\mathrm{rank}(\mathbf{N}^*) = \mathrm{rank}(\mathbf{N})\ .\ \blacksquare$$

**Corollary 5.3**. Let $G, G^*$ be as defined in Proposition 5.2, and $C_b \in \mathcal{B}\backslash\mathcal{B}_2(G)$. Then, $\dim\ker(\mathbf{N}^*) = \dim\ker(\mathbf{N}) - 1$ .

<u>Proof</u>: For any matrix $\mathbf{X} \in \mathbb{R}^{m\times r}$, from the Rank-Nullity Theorem [ref], we know

$$\mathrm{rank}(\mathbf{X}) + \dim\ker(\mathbf{X}) = \dim\mathrm{dom}(\mathbf{X}) = r\ .$$

The two matrices **N** and $\mathbf{N}^*$ have the same number of columns, i.e. $r$ ; if $C_b \in \mathcal{B}\backslash\mathcal{B}_2(G)$, then by virtue of Proposition 5.2, $\mathrm{rank}(\mathbf{N}^*) = \mathrm{rank}(\mathbf{N}) + 1$. It follows that

$$\dim\ker(\mathbf{N}^*) = \dim\ker(\mathbf{N}) - 1\ .\ \blacksquare$$

**Proposition 5.4**. Let $G$ be a network, $C_b \in \mathcal{B}(G)$ and $G^*$ the modified network obtained by injection of $\mathbf{e}_b$ into $G$. Let us denote the deficiency of $G$ and $G^*$ by $\delta$ and $\delta^*$, respectively. Then

$$\delta^* = \delta - 1\,, \qquad (16)$$

if and only if $C_b \in \mathcal{B} \setminus \mathcal{B}_2$. Moreover, $\delta^* = \delta$, if and only if $C_b \in \mathcal{B}_2$.

<u>Proof</u>: From the definition of deficiency,

$$\delta^* = n^* - \ell^* - s^*\ .$$

Note that $G^*$ has the same complexes as $G$, hence, $n^* = n$. Moreover, the two networks have identical linkage structures, hence, $\ell^* = \ell$. Finally, the rank of a CRN is defined as the dimension of the stoichiometric subspace, which is the rank of the stoichiometry matrix; hence, $s^* = \mathrm{rank}(\mathbf{N}^*)$ and $s = \mathrm{rank}(\mathbf{N})$. Therefore,

$$\delta^* = \delta + \mathrm{rank}(\mathbf{N}) - \mathrm{rank}(\mathbf{N}^*)\ .$$

The rest of the proof is trivial, given the statement of Proposition 5.2. $\blacksquare$

**Theorem 6.1**. The maximum length of the sequence $G \to G_1 \to \cdots \to G_d$, that is, $d$ is independent of the choice of nonstoichiometric BCs used to construct it.

<u>Proof</u>: Without loss of generality, let us suppose the balanced complexes iteratively chosen from the set $\mathcal{B} \setminus \mathcal{B}_2 = \left\{C_{b_j}\right\}_{j=1}^{|\mathcal{B}\setminus\mathcal{B}_2|}$ to inject and construct the sequence of modified networks $G \to G_1 \to \cdots \to G_d$ are $C_{b_1}, C_{b_2}, \cdots, C_{b_d}$ , corresponding to vectors $\mathbf{e}_{b_1}, \mathbf{e}_{b_2}, \cdots, \mathbf{e}_{b_d}$ . Suppose

one cannot go further, i.e. no remaining complexes in $\left\{C_{b_j}\right\}_{j=d+1}^{|\mathcal{B}\setminus\mathcal{B}_2|}$ is a nonstoichiometric BC for $G_d$.

Let us now define the matrices

$$\begin{aligned}\mathbf{E} &= \left[\mathbf{e}_{b_1}\ \ \mathbf{e}_{b_2}\ \ \cdots\ \ \mathbf{e}_{b_{|\mathcal{B}\setminus\mathcal{B}_2|}}\right] \ , \\ \mathbf{E}_d &= \left[\mathbf{e}_{b_1}\ \ \mathbf{e}_{b_2}\ \ \cdots\ \ \mathbf{e}_{b_d}\right] \ , \\ \mathbf{E}_{\setminus d} &= \left[\mathbf{e}_{b_{d+1}}\ \ \mathbf{e}_{b_{d+2}}\ \ \cdots\ \ \mathbf{e}_{b_{|\mathcal{B}\setminus\mathcal{B}_2|}}\right] \ .\end{aligned}$$

Given the way the sequence $G \to G_1 \to \cdots \to G_d$ is constructed, it follows from Eq. (S11) that

$$\mathbf{N}_d = \begin{bmatrix}\mathbf{N}_{d-1} \\ \mathbf{e}_{b_d}{}^{\mathrm{T}}\mathbf{A}\end{bmatrix} = \begin{bmatrix}\begin{bmatrix}\mathbf{N}_{d-2} \\ \mathbf{e}_{b_{d-1}}{}^{\mathrm{T}}\mathbf{A}\end{bmatrix} \\ \mathbf{e}_{b_d}{}^{\mathrm{T}}\mathbf{A}\end{bmatrix} = \cdots = \begin{bmatrix}\mathbf{N} \\ \mathbf{E}_d{}^{\mathrm{T}}\mathbf{A}\end{bmatrix} \ .$$

Moreover, from Propositions 5.2 and 5.4, it follows that

$$\begin{aligned}&\mathrm{rank}(\mathbf{N}_d) = \mathrm{rank}(\mathbf{N}) + d \ , \\ &\delta_d \coloneqq \delta(G_d) = \delta - d \ .\end{aligned}$$

Next, we claim that

$$\mathrm{rank}\left(\mathbf{N}_d = \begin{bmatrix}\mathbf{N} \\ \mathbf{E}_d{}^{\mathrm{T}}\mathbf{A}\end{bmatrix}\right) = \mathrm{rank}\left(\begin{bmatrix}\mathbf{N} \\ \mathbf{E}^{\mathrm{T}}\mathbf{A}\end{bmatrix}\right) .$$

We prove this claim by contradiction: Let us assume $\mathrm{rank}(\mathbf{N}_d) \neq \mathrm{rank}\left(\begin{bmatrix}\mathbf{N} \\ \mathbf{E}^{\mathrm{T}}\mathbf{A}\end{bmatrix}\right)$; note that

$$\begin{bmatrix}\mathbf{N} \\ \mathbf{E}^{\mathrm{T}}\mathbf{A}\end{bmatrix} = \begin{bmatrix}\mathbf{N} \\ \mathbf{E}_d{}^{\mathrm{T}}\mathbf{A} \\ \mathbf{E}_{\setminus d}{}^{\mathrm{T}}\mathbf{A}\end{bmatrix} = \begin{bmatrix}\mathbf{N}_d \\ \mathbf{E}_{\setminus d}{}^{\mathrm{T}}\mathbf{A}\end{bmatrix} \ ,$$

hence, the row space of $\mathbf{N}_d$ is a subset of the row space of $\begin{bmatrix}\mathbf{N} \\ \mathbf{E}^{\mathrm{T}}\mathbf{A}\end{bmatrix}$. If the two matrices do not have equal ranks, then there exists a vector $\mathbf{f} \in \left\{\mathbf{e}_{b_{d+1}}, \mathbf{e}_{b_{d+2}}, \cdots, \mathbf{e}_{b_{|\mathcal{B}\setminus\mathcal{B}_2|}}\right\}$, such that

$$\mathrm{rank}\left(\begin{bmatrix}\mathbf{N}_d \\ \mathbf{f}^{\mathrm{T}}\mathbf{A}\end{bmatrix}\right) = \mathrm{rank}(\mathbf{N}_d) + 1 \ .$$

According to Proposition 5.2, this means $\left\{C_{b_j}\right\}_{j=d+1}^{|\mathcal{B}\setminus\mathcal{B}_2|}$ contains a nonstoichiometric BC for $G_d$, which is a contradiction. Therefore, we must have

$$\mathrm{rank}(\mathbf{N}_d) = \mathrm{rank}\left(\begin{bmatrix}\mathbf{N} \\ \mathbf{E}^{\mathrm{T}}\mathbf{A}\end{bmatrix}\right) \ .$$

Hence, the drop in deficiency, $d$, is equal to

$$d = \mathrm{rank}\left(\begin{bmatrix}\mathbf{N} \\ \mathbf{E}^{\mathrm{T}}\mathbf{A}\end{bmatrix}\right) - \mathrm{rank}(\mathbf{N}) \ ,$$

which means $d$ is independent of the order of BCs used to construct the sequence of modified networks.

Even though the sequence $G \to G_1 \to \cdots \to G_d$ itself may depend on the choice of nonstoichiometric BCs injected, its maximum length is a constant. ■

**Theorem 6.2**. Suppose $\mathcal{B} \setminus \mathcal{B}_2 = \left\{C_{b_j}\right\}_{j=1}^{|\mathcal{B}\setminus\mathcal{B}_2|}$ be the set of nonstoichiometric BCs for $G$. Let us construct the matrix $\mathbf{E}_{\mathcal{B}\setminus\mathcal{B}_2}$ as $\mathbf{E}_{\mathcal{B}\setminus\mathcal{B}_2} = \left[\mathbf{e}_{b_1}\ \ \mathbf{e}_{b_2}\ \ \cdots\ \ \mathbf{e}_{b_{|\mathcal{B}\setminus\mathcal{B}_2|}}\right]$. The effective deficiency of $G$ is equal to

$$\delta^{\text{eff}} = \delta + \text{rank}(\mathbf{N}) - \text{rank}\left(\begin{bmatrix}\mathbf{N} \\ \mathbf{E}_{\mathcal{B}\setminus\mathcal{B}_2}{}^{\mathrm{T}}\mathbf{A}\end{bmatrix}\right).$$

<u>Proof</u>: It has already been shown in proof of Theorem 6.1 that

$$d = \text{rank}\left(\begin{bmatrix}\mathbf{N} \\ \mathbf{E}_{\mathcal{B}\setminus\mathcal{B}_2}{}^{\mathrm{T}}\mathbf{A}\end{bmatrix}\right) - \text{rank}(\mathbf{N}) \ ,$$

which, given the definition of $\delta^{\text{eff}}$, readily yields the identity $\delta^{\text{eff}} = \delta + \text{rank}(\mathbf{N}) - \text{rank}\left(\begin{bmatrix}\mathbf{N} \\ \mathbf{E}_{\mathcal{B}\setminus\mathcal{B}_2}{}^{\mathrm{T}}\mathbf{A}\end{bmatrix}\right)$. ■

**Proposition 6.3**. Suppose $\mathcal{B} = \left\{C_{b_j}\right\}_{j=1}^{|\mathcal{B}|}$ be the set of BCs for $G$, and $\mathbf{E}_{\mathcal{B}}$ defined as $\mathbf{E}_{\mathcal{B}} = \left[\mathbf{e}_{b_1}\ \ \mathbf{e}_{b_2}\ \ \cdots\ \ \mathbf{e}_{b_{|\mathcal{B}|}}\right]$. Then, the drop in deficiency, $d$, is the codimension of $\text{im}(\mathbf{Y}^{\mathrm{T}}) + \text{im}(\mathbf{U})$ in $\text{im}(\mathbf{Y}^{\mathrm{T}}) + \text{im}(\mathbf{U}) + \text{im}(\mathbf{E}_{\mathcal{B}})$, that is

$$d = \dim(\text{im}(\mathbf{Y}^{\mathrm{T}}) + \text{im}(\mathbf{U}) + \text{im}(\mathbf{E}_{\mathcal{B}})/\text{im}(\mathbf{Y}^{\mathrm{T}}) + \text{im}(\mathbf{U}))\,,$$

where $V/W$ denotes the quotient space of $V$ by $W$.

<u>Proof</u>: Let us temporarily define the parameter $d_0$ as

$$d_0 \coloneqq \dim(\text{im}(\mathbf{Y}^{\mathrm{T}}) + \text{im}(\mathbf{U}) + \text{im}(\mathbf{E}_{\mathcal{B}})/\text{im}(\mathbf{Y}^{\mathrm{T}}) + \text{im}(\mathbf{U}))\,.$$

It follows immediately from the definition that $d_0$ is equal to

$$d_0 = \text{rank}([\mathbf{Y}^{\mathrm{T}}\quad \mathbf{U}\quad \mathbf{E}_{\mathcal{B}}]) - \text{rank}([\mathbf{Y}^{\mathrm{T}}\quad \mathbf{U}])\,.$$

Given the shared columns of the two matrices and given the definition of rank, this means there exist $d_0$ columns in $\mathbf{E}_{\mathcal{B}}$, say $\{\mathbf{e}_j\}_{j=1}^{d_0}$, which do not lie in the column span of $[\mathbf{Y}^{\mathrm{T}}\quad \mathbf{U}]$, that is

$$d_0 = \text{rank}([\mathbf{Y}^{\mathrm{T}}\quad \mathbf{U}\quad \mathbf{e}_1\,\mathbf{e}_2\cdots\mathbf{e}_{d_0}]) - \text{rank}([\mathbf{Y}^{\mathrm{T}}\quad \mathbf{U}])\,.$$

Furthermore, any other column of $\mathbf{E}_{\mathcal{B}}$ lies in the columns span of $\left[\mathbf{Y}^{\mathrm{T}} \quad \mathbf{U} \quad \mathbf{e}_1 \ \mathbf{e}_2 \ \cdots \ \mathbf{e}_{d_0}\right]$. Obviously, all complexes $\{C_j\}_{j=1}^{d_0}$ are nonstoichiometric BCs for $G$. We claim the vectors $\{\mathbf{e}_j\}_{j=1}^{d_0}$ can be successively injected to $G$, which yields a sequence of modified networks $G \to G_1 \to \cdots \to G_{d_0}$, the deficiencies of which strictly decreases at each iteration. This is easy to prove by contradiction: Suppose injection of $\mathbf{e}_k$ does not decrease the deficiency of $G_{k-1}$, that is, $\mathbf{N}_k$ and $\mathbf{N}_{k-1}$ have the same rank, for some $1 < k \leq d_0$. We have

$$\mathbf{N}_k = \begin{bmatrix} \mathbf{N}_{k-1} \\ \mathbf{e}_k{}^{\mathrm{T}}\mathbf{A} \end{bmatrix}.$$

It follows that $\mathbf{A}^{\mathrm{T}}\mathbf{e}_k = \mathbf{A}^{\mathrm{T}}\mathbf{Y}_{k-1}{}^{\mathrm{T}}\boldsymbol{\eta}$ for some vector $\boldsymbol{\eta}$.

Now $\mathbf{Y}_{k-1}{}^{\mathrm{T}} = [\mathbf{Y}^{\mathrm{T}} \quad \mathbf{e}_1 \ \cdots \ \mathbf{e}_{k-1}]$; we partition $\boldsymbol{\eta} = \begin{bmatrix} \boldsymbol{\eta}_1 \\ \boldsymbol{\eta}_2 \end{bmatrix}$ accordingly to obtain

$$\mathbf{A}^{\mathrm{T}}\mathbf{e}_k = \mathbf{A}^{\mathrm{T}}\mathbf{Y}^{\mathrm{T}}\boldsymbol{\eta}_1 + \mathbf{A}^{\mathrm{T}}[\mathbf{e}_1 \ \cdots \ \mathbf{e}_{k-1}]\boldsymbol{\eta}_2 \,.$$

The matrix $\mathbf{U}$ has been defined to provide a basis for kernel of $\mathbf{A}^{\mathrm{T}}$; therefore, there exist vector $\boldsymbol{\xi}$ such that

$$\mathbf{e}_k = \mathbf{Y}^{\mathrm{T}}\boldsymbol{\eta}_1 + [\mathbf{e}_1 \ \cdots \ \mathbf{e}_{k-1}]\boldsymbol{\eta}_2 + \mathbf{U}\,\boldsymbol{\xi} \,.$$

In this case, one can eliminate the column $\mathbf{e}_k$ from the matrix $\left[\mathbf{Y}^{\mathrm{T}} \quad \mathbf{U} \quad \mathbf{e}_1 \, \mathbf{e}_2 \cdots \mathbf{e}_{d_0}\right]$, without changing its rank. It follows that

$$\begin{aligned} &\operatorname{rank}\left(\left[\mathbf{Y}^{\mathrm{T}} \quad \mathbf{U} \quad \mathbf{e}_1 \, \mathbf{e}_2 \cdots \mathbf{e}_{d_0}\right]\right) = \\ &\operatorname{rank}\left(\left[\mathbf{Y}^{\mathrm{T}} \quad \mathbf{U} \quad \mathbf{e}_1 \, \mathbf{e}_2 \cdots \mathbf{e}_{\mathrm{k}-1} \, \mathbf{e}_{\mathrm{k}+1} \cdots \mathbf{e}_{d_0}\right]\right) < \operatorname{rank}([\mathbf{Y}^{\mathrm{T}} \quad \mathbf{U}]) + d_0 \end{aligned}$$

which is a contradiction.

As a result, $\delta_{d_0} = \delta - d_0$. Given the maximality assumption for $d_0$ and the definition of the effective deficiency, it follows that $d_0 = d$. ■

#### S2 Detailed factorizations for the toy examples

##### S2.1 Toy network I: a type-I nonstoichiometric BC

Let us consider the conversion diagram in Fig. S1.

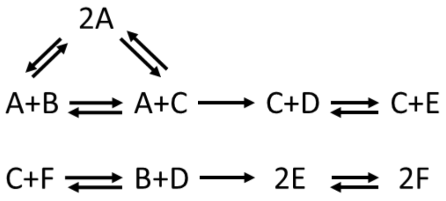


**Fig. S1. A nonstoichiometric BC in a toy network.**
The basic conversion diagram depicts a network operating in a canonical flux regime, with $m = 6$ species, $n = 9$ complexes, and $r = 8$ reactions.

Let us sort the species by alphabetical order and sort the complexes as follows: $(C_1, \cdots, C_9) = (A + B, A + C, C + D, C + E, C + F,\ B + D, 2E, 2F, 2A)$. The stoichiometric map $\mathbf{Y}$ and the incidence matrix $\mathbf{A}$ for this network can then be formed as follows

$$\mathbf{Y} = \begin{bmatrix} 1 & 1 & 0 & 0 & 0 & 0 & 0 & 0 & 2 \\ 1 & 0 & 0 & 0 & 0 & 1 & 0 & 0 & 0 \\ 0 & 1 & 1 & 1 & 1 & 0 & 0 & 0 & 0 \\ 0 & 0 & 1 & 0 & 0 & 1 & 0 & 0 & 0 \\ 0 & 0 & 0 & 1 & 0 & 0 & 2 & 2 & 0 \\ 0 & 0 & 0 & 0 & 1 & 0 & 0 & 0 & 0 \end{bmatrix},$$

$$\mathbf{A} = \begin{bmatrix} 1 & 0 & -1 & 0 & 0 & 0 & 0 & 0 \\ 0 & 1 & 1 & -1 & 0 & 0 & 0 & 0 \\ 0 & 0 & 0 & 1 & -1 & 0 & 0 & 0 \\ 0 & 0 & 0 & 0 & 1 & 0 & 0 & 0 \\ 0 & 0 & 0 & 0 & 0 & -1 & 0 & 0 \\ 0 & 0 & 0 & 0 & 0 & 1 & -1 & 0 \\ 0 & 0 & 0 & 0 & 0 & 0 & 1 & -1 \\ 0 & 0 & 0 & 0 & 0 & 0 & 0 & 1 \\ -1 & -1 & 0 & 0 & 0 & 0 & 0 & 0 \end{bmatrix}.$$

Given the irreversibility pattern in the network and how matrix $\mathbf{A}$ was arranged, it follows that $R_4$ and $R_7$ are the only two irreversible reactions. As mentioned in the article, one can partition $\mathbf{A}$ into blocks $\mathbf{A}^{\mathrm{nl}}$ and $\mathbf{A}^{\mathrm{zl}}$, as follows

$$\mathbf{A}^{\mathrm{nl}} = \begin{bmatrix} 1 & 0 & -1 & 0 & 0 & 0 \\ 0 & 1 & 1 & 0 & 0 & 0 \\ 0 & 0 & 0 & -1 & 0 & 0 \\ 0 & 0 & 0 & 1 & 0 & 0 \\ 0 & 0 & 0 & 0 & -1 & 0 \\ 0 & 0 & 0 & 0 & 1 & 0 \\ 0 & 0 & 0 & 0 & 0 & -1 \\ 0 & 0 & 0 & 0 & 0 & 1 \\ -1 & -1 & 0 & 0 & 0 & 0 \end{bmatrix}, \quad \mathbf{A}^{\mathrm{zl}} = \begin{bmatrix} 0 & 0 \\ -1 & 0 \\ 1 & 0 \\ 0 & 0 \\ 0 & 0 \\ 0 & -1 \\ 0 & 1 \\ 0 & 0 \\ 0 & 0 \end{bmatrix}.$$

Given the linkage structure of this network specified by matrix $\mathbf{A}$, we can form the complex to linkage-class incidence matrix $\mathbf{U}$ as follows

$$\mathbf{U} = \begin{bmatrix} 1 & 0 \\ 1 & 0 \\ 1 & 0 \\ 1 & 0 \\ 0 & 1 \\ 0 & 1 \\ 0 & 1 \\ 0 & 1 \\ 1 & 0 \end{bmatrix}.$$

Clearly, the columns of this matrix form a basis for the left nullspace of $\mathbf{A}$.

It can be shown that the complex $(2A)$ (indexed by vector $\mathbf{e}_9$) has a type-I nonstoichiometric factorization with the following parameters

$$\boldsymbol{\zeta}_1 = \begin{bmatrix} 1 \\ 0 \\ 0 \\ 0 \\ 0 \\ 0 \end{bmatrix}, \boldsymbol{\xi}_1 = \begin{bmatrix} -1 \\ 0 \end{bmatrix}, \boldsymbol{\theta}_1 = \begin{bmatrix} 0 \\ 0 \\ 1 \\ 1 \\ 0 \\ 0 \\ 0 \\ 0 \\ 0 \end{bmatrix}, \qquad \boldsymbol{\zeta}_2 = \begin{bmatrix} 1 \\ 0 \\ 0 \\ 1 \\ 1 \\ 1 \end{bmatrix}, \boldsymbol{\xi}_2 = \begin{bmatrix} -1 \\ -1 \end{bmatrix}, \boldsymbol{\theta}_2 = \begin{bmatrix} 0 \\ 0 \\ 0 \\ 0 \\ 0 \\ 0 \\ 1 \\ 1 \\ 0 \end{bmatrix};$$

that is,

$$\begin{cases} \mathbf{e}_9 = \mathbf{Y}^{\mathrm{T}}\boldsymbol{\zeta}_1 + \mathbf{U}\,\boldsymbol{\xi}_1 + \boldsymbol{\theta}_1 \\ \mathbf{e}_9 = \mathbf{Y}^{\mathrm{T}}\boldsymbol{\zeta}_2 + \mathbf{U}\,\boldsymbol{\xi}_2 - \boldsymbol{\theta}_2 \\ {\mathbf{A}^{\mathrm{nl}}}^{\mathrm{T}}\,[\boldsymbol{\theta}_1 \;\; \boldsymbol{\theta}_2] = \mathbf{0} \\ {\mathbf{A}^{\mathrm{zl}}}^{\mathrm{T}}\,[\boldsymbol{\theta}_1 \;\; \boldsymbol{\theta}_2] \geqslant \mathbf{0} \end{cases}$$

It follows that $C_9$ is a type-I nonstoichiometric BC for the network. It is also easy to show that no other complex is balanced in this network. The network is of deficiency $\delta = n - \ell - s = 9 - 2 - 5 = 2$. However, since vector $\mathbf{e}_9$ represents a nonstoichiometric BC, one can inject $\mathbf{e}_9$ into the network to obtain an equivalent network of lower deficiency. Moreover, the obtained network will have no nonstoichiometric BCs, that is, $d = 1$. It follows that $\delta^{\mathrm{eff}} = \delta - d = 1$.

In addition, using very simple tools such as LP solvers, one may easily show that the maximum flux going through irreversible reactions $R_4$ and $R_7$ is zero, i.e. they are both blocked at steady state. This is consistent with the prediction of Proposition 4.1.

Now, suppose we remove the two blocked reactions, which yields a reduced network shown in Fig. S2.

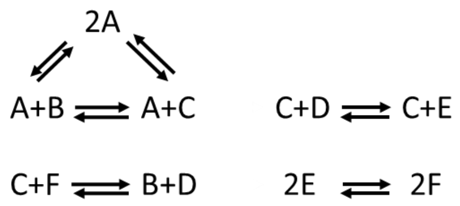


**Fig. S2. A nonstoichiometric BC in a toy network.**
The reduced conversion diagram depicts a network operating in a canonical flux regime, with $m = 6$ species, $n = 9$ complexes, and $r = 6$ reactions.

One may associate this network with the following matrices $\mathbf{Y}'$ and $\mathbf{A}'$.

$$\mathbf{Y}' = \mathbf{Y} = \begin{bmatrix} 1 & 1 & 0 & 0 & 0 & 0 & 0 & 0 & 2 \\ 1 & 0 & 0 & 0 & 0 & 1 & 0 & 0 & 0 \\ 0 & 1 & 1 & 1 & 1 & 0 & 0 & 0 & 0 \\ 0 & 0 & 1 & 0 & 0 & 1 & 0 & 0 & 0 \\ 0 & 0 & 0 & 1 & 0 & 0 & 2 & 2 & 0 \\ 0 & 0 & 0 & 0 & 1 & 0 & 0 & 0 & 0 \end{bmatrix},$$

$$\mathbf{A}' = \mathbf{A}^{\mathrm{nl}} = \begin{bmatrix} 1 & 0 & -1 & 0 & 0 & 0 \\ 0 & 1 & 1 & 0 & 0 & 0 \\ 0 & 0 & 0 & -1 & 0 & 0 \\ 0 & 0 & 0 & 1 & 0 & 0 \\ 0 & 0 & 0 & 0 & -1 & 0 \\ 0 & 0 & 0 & 0 & 1 & 0 \\ 0 & 0 & 0 & 0 & 0 & -1 \\ 0 & 0 & 0 & 0 & 0 & 1 \\ -1 & -1 & 0 & 0 & 0 & 0 \end{bmatrix}.$$

Note that removing the two blocked reactions would alter the linkage structure of the network, hence the matrix $\mathbf{U}'$ is expanded as follows

$$\mathbf{U}' = \begin{bmatrix} 1 & 0 & 0 & 0 \\ 1 & 0 & 0 & 0 \\ 0 & 1 & 0 & 0 \\ 0 & 1 & 0 & 0 \\ 0 & 0 & 1 & 0 \\ 0 & 0 & 1 & 0 \\ 0 & 0 & 0 & 1 \\ 0 & 0 & 0 & 1 \\ 1 & 0 & 0 & 0 \end{bmatrix}.$$

As a result, the complex $C_9 = 2A$ has turned into a stoichiometric BC for the reduced network, with the following stoichiometric factorization

$$\mathbf{e}_9 = \mathbf{Y}'^{\mathrm{T}}\boldsymbol{\zeta} + \mathbf{U}'\,\boldsymbol{\xi}\ , \qquad \boldsymbol{\zeta} = \begin{bmatrix} 1 \\ 0 \\ 0 \\ 0 \\ 0 \\ 0 \end{bmatrix}, \qquad \boldsymbol{\xi} = \begin{bmatrix} -1 \\ 0 \\ 0 \\ 0 \end{bmatrix}$$

Note that for the reduced network, one may calculate the deficiency as

$$\delta' = n' - \ell' - s' = 9 - 4 - 4 = 1\,,$$

which does not correspond to the nominal deficiency $\delta$ calculated above, but actually to the effective deficiency $\delta^{\text{eff}} = 1$.

It is perhaps worth mentioning that changing the irreversibility patterns elsewhere in the original network does not affect the balancing property of $C_9$; however, $C_9$ would not be balanced anymore if one modifies either of $R_4$ and $R_7$. It is precisely the irreversibility of these two reactions in this particular constellation that makes $C_9$ a balanced complex.

##### S2.2 Toy network II: a type-II nonstoichiometric BC

Let us next consider the following toy network

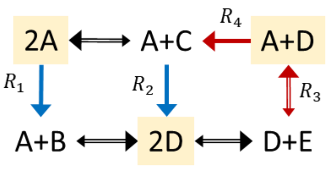


**Fig. S3. Type-II nonstoichiometric BCs in a toy network.**
The conversion diagram portrays a network with $m = 5$ species, $n = 6$ complexes, and $r = 7$ reactions. The network operates in a non-canonical and bounded flux regime, where the fluxes of irreversible reactions $R_1$ and $R_2$ have strictly positive lower bounds ($v_{\text{l},1} = v_{\text{l},2} = 100$), while the fluxes of reactions $R_3$ and $R_4$ have finite upper bounds ($v_{\text{u},3} = v_{\text{u},4} = 200$). The network contains three type-II nonstoichiometric BCs, highlighted in yellow. The rest of complexes are stoichiometric BCs.
For the reversible reaction, the larger arrow size depict the direction of the flux associated with a positive sign.

Let us sort the species by alphabetical order and sort the complexes as follows: $(C_1, \cdots, C_6) = (2A, A + C, A + D, A + B, 2D, D + E)$. The stoichiometric map **Y** and the complex-reaction incidence matrix **A** for this network can then be formed as follows

$$\mathbf{Y} = \begin{bmatrix} 2 & 1 & 1 & 1 & 0 & 0 \\ 0 & 0 & 0 & 1 & 0 & 0 \\ 0 & 1 & 0 & 0 & 0 & 0 \\ 0 & 0 & 1 & 0 & 2 & 1 \\ 0 & 0 & 0 & 0 & 0 & 1 \end{bmatrix},$$

$$\mathbf{A} = \begin{bmatrix} -1 & 0 & 0 & 0 & 1 & 0 & 0 \\ 0 & -1 & 0 & 0 & -1 & 0 & 0 \\ 0 & 0 & 1 & -1 & 0 & 0 & 0 \\ 1 & 0 & 0 & 1 & 0 & -1 & 0 \\ 0 & 1 & 0 & 0 & 0 & 1 & -1 \\ 0 & 0 & -1 & 0 & 0 & 0 & 1 \end{bmatrix}$$

The lower- and upper bounds on flux through reactions $(R_{!}, \cdots, R_7)$ are assigned as follows

$$\mathbf{v}_\text{l} = [100 \quad 100 \quad -300 \quad 0 \quad -300 \quad -300 \quad -300]^\text{T}\,,$$

$$\mathbf{v}_\text{u} = [300 \quad 300 \quad 200 \quad 200 \quad 300 \quad 300 \quad 300]^\text{T}\,.$$

The complexes $\mathbf{e}_2, \mathbf{e}_4$ and $\mathbf{e}_6$ are trivial BCs (strictly-stoichiometric) with the following factorizations:

$$\mathbf{e}_2 = \mathbf{Y}^{\mathrm{T}}\begin{bmatrix}0\\0\\1\\0\\0\end{bmatrix}, \qquad \mathbf{e}_4 = \mathbf{Y}^{\mathrm{T}}\begin{bmatrix}0\\1\\0\\0\\0\end{bmatrix}, \qquad \mathbf{e}_6 = \mathbf{Y}^{\mathrm{T}}\begin{bmatrix}0\\0\\0\\0\\1\end{bmatrix}.$$

Next, let us take a complex, say $e_5 \leftrightarrow (2D)$, that is not stoichiometrically balanced. It is not difficult to show that the variables

$$\boldsymbol{\zeta}_1 = \begin{bmatrix}0\\-1\\0\\0\\-1\end{bmatrix}, \boldsymbol{\lambda}_{\mathrm{l}1} = \begin{bmatrix}1\\1\\0\\0\\0\\0\\0\end{bmatrix}, \boldsymbol{\lambda}_{\mathrm{u}1} = \begin{bmatrix}0\\0\\1\\0\\0\\0\\0\end{bmatrix}, \qquad \boldsymbol{\zeta}_2 = \begin{bmatrix}0\\1\\0\\1\\0\end{bmatrix}, \boldsymbol{\lambda}_{\mathrm{l}2} = \begin{bmatrix}1\\1\\0\\0\\0\\0\\0\end{bmatrix}, \boldsymbol{\lambda}_{\mathrm{u}2} = \begin{bmatrix}0\\0\\0\\1\\0\\0\\0\end{bmatrix}$$

satisfy the following nonstoichiometric factorization for $\mathbf{e}_5$.

$$\begin{cases}\mathbf{A}^{\mathrm{T}}\mathbf{e}_5 = \mathbf{A}^{\mathrm{T}}\mathbf{Y}^{\mathrm{T}}\boldsymbol{\zeta}_1 + \boldsymbol{\lambda}_{\mathrm{l}1} - \boldsymbol{\lambda}_{\mathrm{u}1}\\ \mathbf{A}^{\mathrm{T}}\mathbf{e}_5 = \mathbf{A}^{\mathrm{T}}\mathbf{Y}^{\mathrm{T}}\boldsymbol{\zeta}_2 + \boldsymbol{\lambda}_{\mathrm{u}2} - \boldsymbol{\lambda}_{\mathrm{l}2}\\ \mathbf{v}_{\mathrm{l}}^{\mathrm{T}}\boldsymbol{\lambda}_{\mathrm{l}1} - \mathbf{v}_{\mathrm{u}}^{\mathrm{T}}\boldsymbol{\lambda}_{\mathrm{u}1} = \mathbf{v}_{\mathrm{l}}^{\mathrm{T}}\boldsymbol{\lambda}_{\mathrm{l}2} - \mathbf{v}_{\mathrm{u}}^{\mathrm{T}}\boldsymbol{\lambda}_{\mathrm{u}2} = 0\ , \quad t = 1{,}2\\ \boldsymbol{\lambda}_{\mathrm{l}1}, \boldsymbol{\lambda}_{\mathrm{l}2}, \boldsymbol{\lambda}_{\mathrm{u}1}, \boldsymbol{\lambda}_{\mathrm{u}2} \geqslant \mathbf{0}\end{cases}$$
